# TRAF4 inhibits bladder cancer progression by promoting BMP/SMAD signalling pathway

**DOI:** 10.1101/2020.10.12.335588

**Authors:** Prasanna Vasudevan Iyengar, Dieuwke Louise Marvin, Dilraj Lama, Tuan Zea Tan, Sudha Suriyamurthy, Feng Xie, Maarten van Dinther, Hailiang Mei, Chandra Shekhar Verma, Long Zhang, Laila Ritsma, Peter ten Dijke

**Affiliations:** Department of Cell and Chemical Biology, Leiden University Medical Center, Leiden 2333ZC, The Netherlands; Oncode Institute, The Netherlands; Department of Microbiology, Tumor and Cell Biology, Karolinska Institutet, Biomedicum Quarter 7B-C Solnavagen 9, 17165 Solna, Stockholm, Sweden; Bioinformatics Institute (A*STAR), 30 Biopolis street, 07-01 Matrix, Singapore 138671; Cancer Science Institute of Singapore, National University of Singapore, Singapore 117599; Life Sciences Institute, Zhejiang University, Hangzhou, Zhejiang 310058, China; Institutes of Biology and Medical Science, Soochow University, Suzhou 215123, China; Sequencing Analysis Support Core, Department of Biomedical Data Sciences, Leiden University Medical Center, 2333 ZC, Leiden, The Netherlands; Department of Biological sciences, National University of Singapore, 14 Science Drive 4, Singapore 117543; School of Biological sciences, Nanyang Technological University, 50 Nanyang Drive, Singapore 637551

**Keywords:** Bladder cancer, Epithelial to mesenchymal transition, bone morphogenetic protein, NF-κB, E3 ubiquitin ligases

## Abstract

Bladder cancer is one of the most prevalent cancer types in the world, frequently exhibiting invasion and metastasis and therefore associated with poor prognosis. It is a progressive disease with high recurrence rates even after surgery, which calls for the urgent need for early intervention and diagnosis. The E3 ubiquitin ligase TNF Receptor Associated Factor 4 (TRAF4) has been largely implicated as a tumour-promoting element in a wide range of cancers. Over-expression and amplification of this gene product has been a common observation in breast and other metastatic tumours. In contrast, we observed that expression of TRAF4 negatively correlated with overall patient survival. Moreover, its expression was repressed at epigenetic levels in aggressive bladder cancer cells. We also describe an ERK kinase phosphorylation site on TRAF4 that inhibits its stability and localization to plasma membrane in such cells. Furthermore, knockdown of TRAF4 in epithelial bladder cancer cell lines leads to gain of mesenchymal genes and a loss of epithelial integrity. Reciprocally, stable over-expression of TRAF4 in mesenchymal cells leads to decreased migratory and invasive properties. Transcriptomic analysis upon TRAF4 mis-expression in bladder cancer cell lines revealed that higher TRAF4 expression enhanced BMP/SMAD and dampened NF-κB signalling pathways. Importantly, this notion was confirmed in bladder cancer patient material. Mechanistically, we showed that TRAF4 targets the E3 ubiquitin ligase SMURF1, a negative regulator of BMP/SMAD signalling, for proteasomal degradation in bladder cancer cells. We show that genetic and pharmacological inhibition of SMURF1 or its function inhibited migration of aggressive (mesenchymal) bladder cancer cells.

## Introduction

Metastasis is the key determinant to poor cancer patient survival outcome. Hence, numerous studies in the past decade have focused on thwarting the metastatic potential of cancer cells to allow for early diagnosis and therapeutic intervention in a manner that is optimal for each patient. Epithelial-Mesenchymal Transition or EMT has received a lot of attention with respect to metastasis of cancer cells. EMT was initially discovered as a critical event during embryogenesis especially during gastrulation and organogenesis[1]; but later studies have revealed the process to have a role during cancer metastasis[2]. During EMT, epithelial cells which generally have higher levels of E-cadherin at their cell-cell junction lose its expression and gain expression of N-cadherin[3]; such events are carried out by an orchestra of transcription factors, the major ones being SNAIL and SLUG. It is important to note however, that EMT is a transient event; during the rapid outgrowth of micro-metastases the reversal of EMT, mesenchymal to epithelial transition is needed. In addition, cancer cells often do not undergo a full transition to mesenchymal state and retain epithelial properties. Importantly, the acquisition of these partial EMT states was found to associate with enhanced metastatic potential[4, 5].

Previously, we have described a RING domain containing E3 ubiquitin ligase TRAF4 as a promoter of metastatic breast cancer[6]. We showed that high TRAF4 expression strongly correlated with poor clinical outcomes for breast cancer patients. Importantly, TRAF4 acts as a catalyst to enhance TGF-β/SMAD and non-SMAD mediated EMT. Several other studies have also shown that TRAF4 is a critical factor for driving prostate, lung, glioma tumour progression[7–10]. We previously demonstrated the involvement of EMT in bladder cancer progression[11]. Bladder cancer is a highly prevalent cancer with poor clinical outcomes; Especially in advanced stages of progression when the cancer starts invading the bladder muscle. We investigated if TRAF4 plays a role in bladder cancer progression given its notorious involvement in EMT and metastatic diseases. Unexpectantly, we found that TRAF expression is positively associated with good prognosis in bladder cancer. We uncovered how TRAF4 expression and/or function become compromised, and elucidated how this triggers an increase in migration, invasion and EMT of bladder cancer cells. Moreover, using transcriptional profiling, genetic and pharmacological intervention approaches we show an involvement of specific pathways. We additionally demonstrate a correlation of TRAF4 expression with downstream signalling components of BMP/SMAD and NF-κB signalling pathways using bladder cancer patient material. Our findings may, therefore, be of importance for therapy of bladder cancer patients with low TRAF4 expression.

## Results

### TRAF4 expression negatively correlates with bladder cancer progression

We investigated the correlation between *TRAF4* mRNA expression and overall survival of bladder cancer patients across all stages. Interestingly, a Kaplan-Meier plot using publicly available TCGA BLCA data obtained from the Human Protein Atlas www.proteinatlas.org revealed that bladder cancer patients with a lower level of *TRAF4* expression had significantly lower survival probability than those with higher *TRAF4* expression (Fig 1A)[12]. To further confirm if these observations also hold true for TRAF4 expression at the protein level, we performed immunohistochemistry using tissue microarray samples obtained from Biomax U.S (BL802b). Our data revealed significant differences in TRAF4 expression between stage 1 tumour samples and stage 2/3, and expression differences between stage 1 with adjacent normal tissue or normal bladder tissue samples were non-significant (Fig 1B and 1C). To further corroborate our initial observations, we cross-checked *TRAF4* expression in a recently compiled meta-cohort study of 2411 bladder tumour data[13]. The classification includes six distinct molecular subtypes: Her2L (Her2-like), Pap (Papillary), Neural (Neu), Lum (Luminal), SCC (Squamous cell carcinoma) and Mes (Mesenchymal). We observed that *TRAF4* expression was the lowest in SCC and Mes subtypes that have the poorest survival outcomes (Fig 1D). EMT scoring can be performed using a specific EMT gene signature[14]. It is noteworthy that SCC and Mes subtypes have the highest EMT scores, meaning that these cancer cells are most likely to be mesenchymal-like in phenotype[13]. Furthermore, we extended our EMT scoring to bladder cancer cell lines. Using publicly available data performed on 59 bladder cancer (human) cell lines [15], we calculated the EMT scores as shown (Fig 1E, Sup. Table 1). We defined cell lines with a negative EMT score as more epithelial-like, whereas cell lines with a positive EMT score as more mesenchymal-like. Consistent with patient data, a significant negative correlation was found between *TRAF4* expression against the EMT scores of 59 bladder cancer cells (Fig 1F). We selected 5 bladder cancer cell lines, RT4 and HT1376 cell lines with negative EMT scores and T24, J82 and UMUC3 with positive EMT scores for further consideration in our study. We next examined if TRAF4 expression correlated with the ‘EMT-status’ of these cell lines at the protein level. As seen in Fig 1G, TRAF4 expression was higher in epithelial cell lines exhibiting higher levels of E-cadherin and lower levels of mesenchymal markers N-cadherin and Vimentin [4]. On the contrary, TRAF4 expression was lower in mesenchymal cell lines with lower E-cadherin levels, but higher N-cadherin and Vimentin levels. Importantly, epithelial cell lines also had higher *TRAF4* mRNA expression levels compared to mesenchymal cell lines (Fig 1H). Collectively, our results suggest that TRAF4 expression is indeed higher in less aggressive epithelial bladder cancer cells as compared to more aggressive mesenchymal bladder cancer cells.

**Figure 1.**
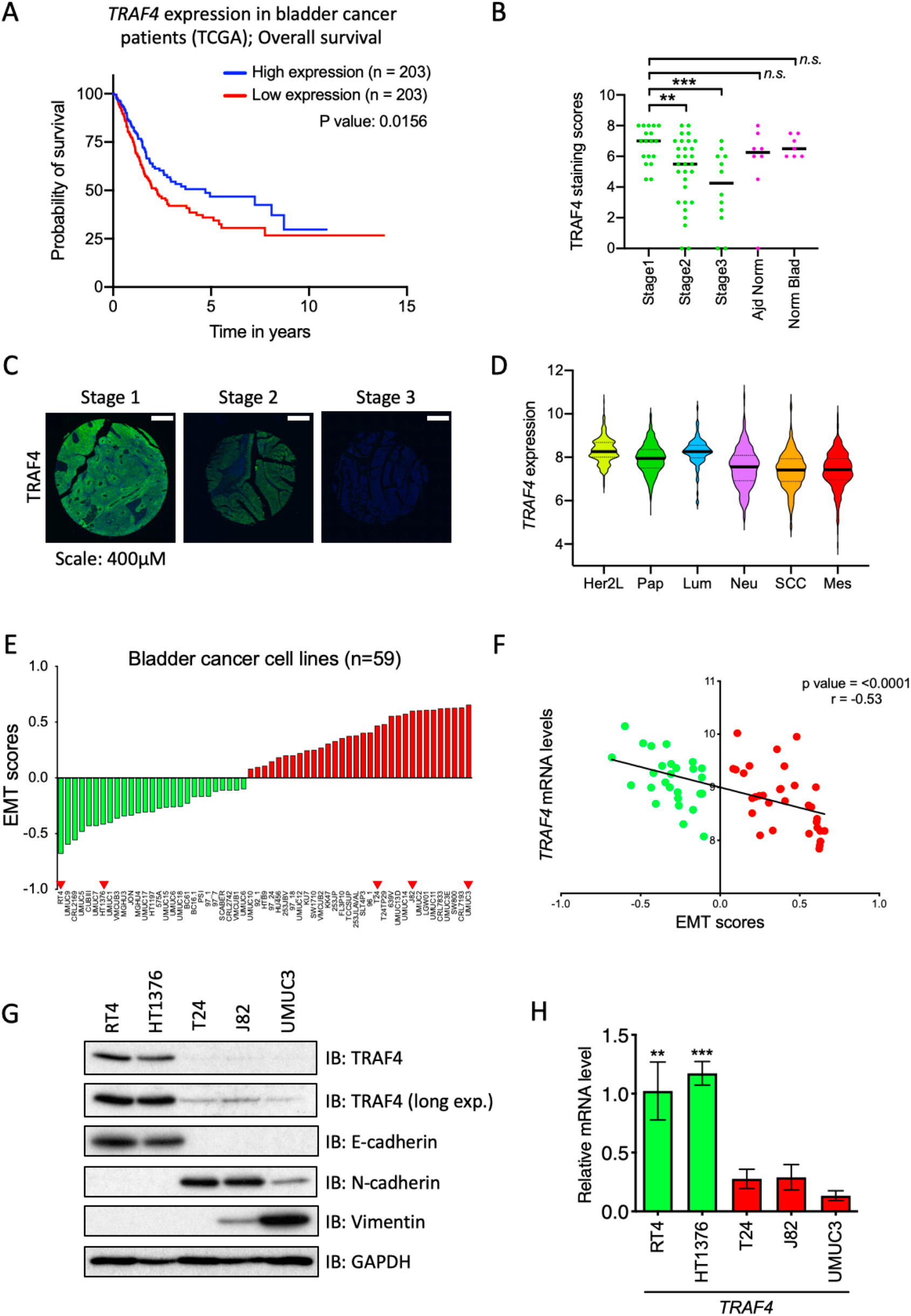
TRAF4 is downregulated in aggressive bladder tumours and mesenchymal bladder cancer cell lines. **A** Kaplan-Meier plot showing overall survival of bladder cancer patients with differential *TRAF4* expression, data obtained and reproduced from TCGA (The Human Protein Atlas), the median FKPM value was taken as *TRAF4* expression cut-off. **B** Graph representing TRAF4 expression through scores obtained from immunohistochemistry of tissue microarray, \*\**P* ≤ 0.01, \*\*\**P* ≤ 0.001 calculated using two-tailed student’s t test; *n.s.* indicates non-significant P value. **C** Representative images of immunohistochemistry performed for TRAF4 expression on tissue microarray from stages 1-3 of bladder tumours. **D** Violin plot shows *TRAF4* expression levels (and distribution) across different subtypes of bladder cancer, Her2L: Her2-like (n=253), Pap: Papillary (n=674), Lum: Luminal (n=107), Neu: Neural (n=448), SCC: Squamous cell carcinoma (n=333) and Mes: Mesenchymal (n=308). Black bars in the middle of the distribution represents Median. The subtypes have been arranged according to their EMT scores. **E** Plot represents EMT scores in 59 bladder cancer cell lines, green bars indicate cell lines with negative EMT scores, red bars indicate cell lines with positive EMT scores; red arrowheads indicate cell lines taken for further investigation. **F** Regression plot for *TRAF4* expression levels vs EMT scores in 59 bladder cancer cell lines. **G** Immunoblot analysis showing TRAF4 and other EMT markers protein expression. **H** Real-time PCR showing *TRAF4* mRNA expression in cell lines. Error bars represent ± SD. Epithelial cell lines (green bars) have significantly higher *TRAF4* expression than mesenchymal cell lines (red bars), \*\**P* ≤ 0.01, \*\*\**P* ≤ 0.001 calculated using two-tailed student’s t test.

### TRAF4 is epigenetically repressed and gets phosphorylated at Serine 334 by ERK kinase

We then sought to determine the reasons behind the low expression of *TRAF4* in mesenchymal cells. We subjected three mesenchymal cell lines to 5’-Azacytidine (a compound that blocks DNA methylation) treatment for a week. As can be seen from Fig 2A, *TRAF4* expression was rescued upon treatment in the cell lines, suggesting that indeed *TRAF4* is epigenetically repressed. We also used a positive control for our experiment, *CDH1* (encoding E-cadherin), which is known to be epigenetically repressed in many mesenchymal cancer cells. Fig 2B shows that treatment with 5’-Azacytidine rescued *CDH1* expression. Surprisingly however, the treatment did not show up-regulation of TRAF4 protein levels, although E-cadherin levels were up-regulated in 2 out of 3 cell lines (Sup. Fig 1A). This suggested that there are additional mechanisms that control TRAF4 protein levels. We then performed a cycloheximide chase assay to determine protein stability. We found that indeed, TRAF4 protein levels are less stable in UMUC3 compared to RT4 and HT1376 (Fig. 1C). To determine if specific post-translational modifications exist that regulate its protein stability, we performed mass-spectrometric analysis, whereby we over-expressed Flag-TRAF4 in 293T cells (Sup. Table 2). We observed that TRAF4 is subjected to several phosphorylation events aimed at its Serine and Threonine residues scattered through its length (Fig 2D).

**Figure 2.**
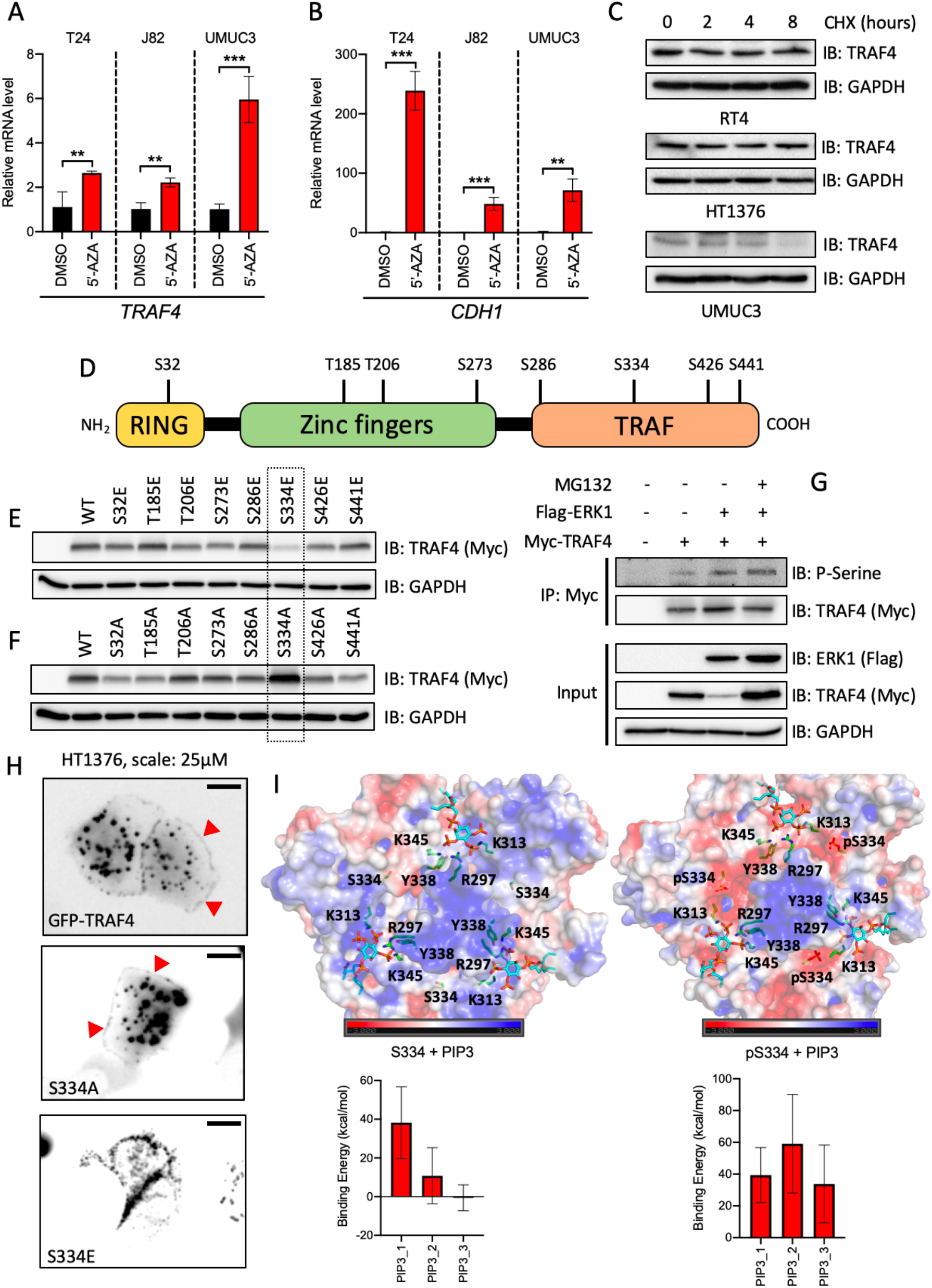
TRAF4 is repressed in mesenchymal (bladder cancer) cell lines at epigenetic and proteomic levels. **A** Real-time PCR result showing changes in *TRAF4* mRNA levels in mesenchymal cell lines after treatment with 5’-Azacitydine; error bars represent ± SD, \*\**P* ≤ 0.01, \*\*\**P* ≤ 0.001 calculated using two-tailed student’s t test. **B** Real-time PCR result showing changes in *CDH1* mRNA levels in cell lines after treatment with 5’-Azacitydine; error bars represent ± SD, \*\**P* ≤ 0.01, \*\*\**P* ≤ 0.001 calculated using two-tailed student’s t test. **C** Immunoblot results showing endogenous TRAF4 levels after treatment with cycloheximide (CHX) using the indicated cell lines. **D** Schematic representation of TRAF4 showing distinct domain structures and candidate phosphorylated serine and threonine sites derived from mass spectrometric analysis. **E** Immunoblot result from 293T transfected with expression constructs for either TRAF4 or TRAF4 glutamic acid mutants. **F** Immunoblot result from 293T transfected with expression constructs for either TRAF4 or TRAF4 alanine mutants. **G** Immunoprecipitation result from 293T transfected with the indicated plasmids and immunoprecipitated with Myc antibody. **H** Immunofluorescence images of HT1376 transfected with expression constructs for either GFP-TRAF4 or GFP-S334 mutants, red arrowheads indicate localization of GFP-TRAF4 and GFP-S334A at the plasma membrane. **I** Representative structures from molecular dynamics simulations of the TRAF domain trimer from TRAF4 in complex with PIP3-dic4 lipid molecules in the unphosphorylated and phosphorylated states of S334. The TRAF domain is shown in electrostatic surface representation (created using the APBS plugin through the Pymol molecular visualization software, Schrondinger) and the color gradient from blue to red indicates the range of electrostatic surface potential kT/e values from strongly positive (+3.0) to strongly negative (−3.0). The residues that are involved in specifically interacting with the lipid headgroup along with the lipid are explicity shown in stick representation and labelled. The lower graphs indicates the binding energies of the individual lipids for the respective systems; the energies were computed using the Molecular Mechanics/Generalized Born Surface Area (MM/GBSA) method by following the same procedure and parameters as described previously[47].

To determine if phosphorylation at these sites affect TRAF4 stability, we performed site-directed mutagenesis to mutate the candidate Serine/Threonine residues to Glutamic acids (E) to mimic, or Alanine residues (A) to block phosphorylation. As can be seen in Fig 2E and 2F, mutating Serine334 to Glutamic acid significantly reduced its stability, and reciprocally, Serine334 to Alanine rescued TRAF4 stability compared to modifications at the other sites. Serine334 site appears to be highly conserved among TRAF4 in other species examined (Sup. Fig 1B). A cycloheximide chase assay revealed that indeed S334E is less stable than WT or S334A mutant (Sup. Fig 1C and 1D). Moreover, addition of proteasomal inhibitor MG132 rescued the instability of S334E mutant suggesting that its unstable due to ubiquitination mediated degradation (Sup. Fig 1E). A phosphorylation prediction tool (Human Protein Reference Database, Phosphomotif finder) revealed that Serine334 could be a putative ERK MAP kinase phosphorylation site. When we over-expressed ERK1 along with TRAF4 in 293T cells, its stability was significantly reduced. Importantly, ERK1 was not able to influence the stability of the S334A mutant (Sup. Fig 1F). MG132 was also able to rescue ERK1 mediated instability of TRAF4 suggesting that ERK1 indeed influences TRAF4’s stability (Sup. Fig 1G). To actually determine if ERK1 can induce phosphorylation of TRAF4 and if this can be enhanced by MG132, we performed an *in vivo* kinase assay. As can be seen from Fig. 2G, co-transfection of ERK1 with TRAF4 indeed increased its phosphorylation and this effect can be further enhanced upon treatment with MG132. Reciprocally, we then assessed if a MEK inhibitor (MEKi, PD0325901) could effectively enhance TRAF4 stability both in over-expressed conditions as well as endogenous levels in mesenchymal cells. Sup. Fig 1H shows that indeed MEKi could slightly enhance levels of TRAF4. The observation also held true in UMUC3 and T24 cell lines, where endogenous TRAF4 levels were stabilized slightly by treatment with MEKi, but TRAF4 mRNA levels remained relatively unaffected (Sup. Fig 1I, 1J, 1K and 1L).

TRAF4 has a ‘TRAF’ domain at its C-terminal region, which mediates trimer formation with other TRAF4 molecules; trimers are capable of binding phosphatidylinositol trisphosphate (PIP3) molecules which tethers them to the plasma membrane[16]. This feature allows TRAF4 molecules to localize to the plasma membrane and regulate tight junction stability[17, 18]. Since, Serine334 is located on the TRAF domain, we hypothesized that phosphorylation of this residue would cause mis-localization of TRAF4. Results in Fig 2H demonstrate that in HT1376, the WT-TRAF4 and S334A mutants localize to cytoplasmic vesicles as well as to the plasma membrane. However, the S334E mutant had a different cytoplasmic localization pattern and more importantly, plasma membrane localization was not observed (also Sup. Fig 2A and 2B). To gain further molecular insights into the effect of S334 phosphorylation on PIP3 binding, we performed modelling and simulations of TRAF domain trimer docked with PIP3 molecules. The modelled complex indicated that S334 is located in the vicinity of the PIP3 headgroup binding site on the TRAF domain (Fig 2I). Phosphorylation of S334 (pS334) distinctly changes the local chemical environment on the TRAF surface by inducing a negative potential (Sup. Fig 2C, 2D, 2E and 2F). This would be electrostatically detrimental for the negatively charged phosphates on the lipid headgroup which is reflected in the positive binding energies of the PIP3 molecules computed from simulations (Fig 2I, Sup. Fig 2G and 2H). The lipid molecules in the S334 phosphorylated state are characterized by unfavourable energetics as compared to S334 in the unphosphorylated state. Taken together, our results suggest that phosphorylation of Serine 334 of TRAF4 by ERK kinase leads to its destabilization via proteasome mediated degradation. Moreover, our modelling predicts that it disrupts the ability to bind PIP3, which in turn interferes with its ability to localize to the plasma membrane.

### Knockdown of TRAF4 in epithelial cell lines leads to loss of epithelial integrity and gain of mesenchymal markers

Since we observed that the expression of TRAF4 is reduced in mesenchymal cells when compared to epithelial bladder cancer cells, we went on to investigate the consequences of TRAF4 knockdown in epithelial cancer cell lines, RT4 and HT1376. Interestingly, depletion of TRAF4 in RT4 cells using two independent short-hairpin RNAs (#4 and #5 with highest knockdown efficiency (Sup. Fig 3A), led to an increase in mRNA expression levels of mesenchymal markers *CDH2* (encoding N-cadherin) and *SNAI2* (encoding SLUG) (Fig 3A). SLUG is a well-studied EMT-transcription factor that has been described to play roles in cadherin switching and malignancy in bladder cancer progression[19]. Its noteworthy that another major EMT-TF, *SNAI1* (encoding SNAIL) consistently gets down-regulated upon TRAF4 knockdown. The increase in SLUG and N-cadherin (but not E-cadherin) expression was also observed at protein levels (Fig 3B). Next, we observed phenotypic changes in the integrities and architecture of cell colonies upon TRAF4 knockdown (Fig 3C). A membrane staining dye revealed that the borders of cells within RT4 cell colonies get disorganized and appearance of loosely attached cells upon TRAF4 knockdown. The membrane staining of colonies in control cells (pLKO vector) resembled staining in RT4 wildtype cells (Sup. Fig 3B).

**Figure 3.**
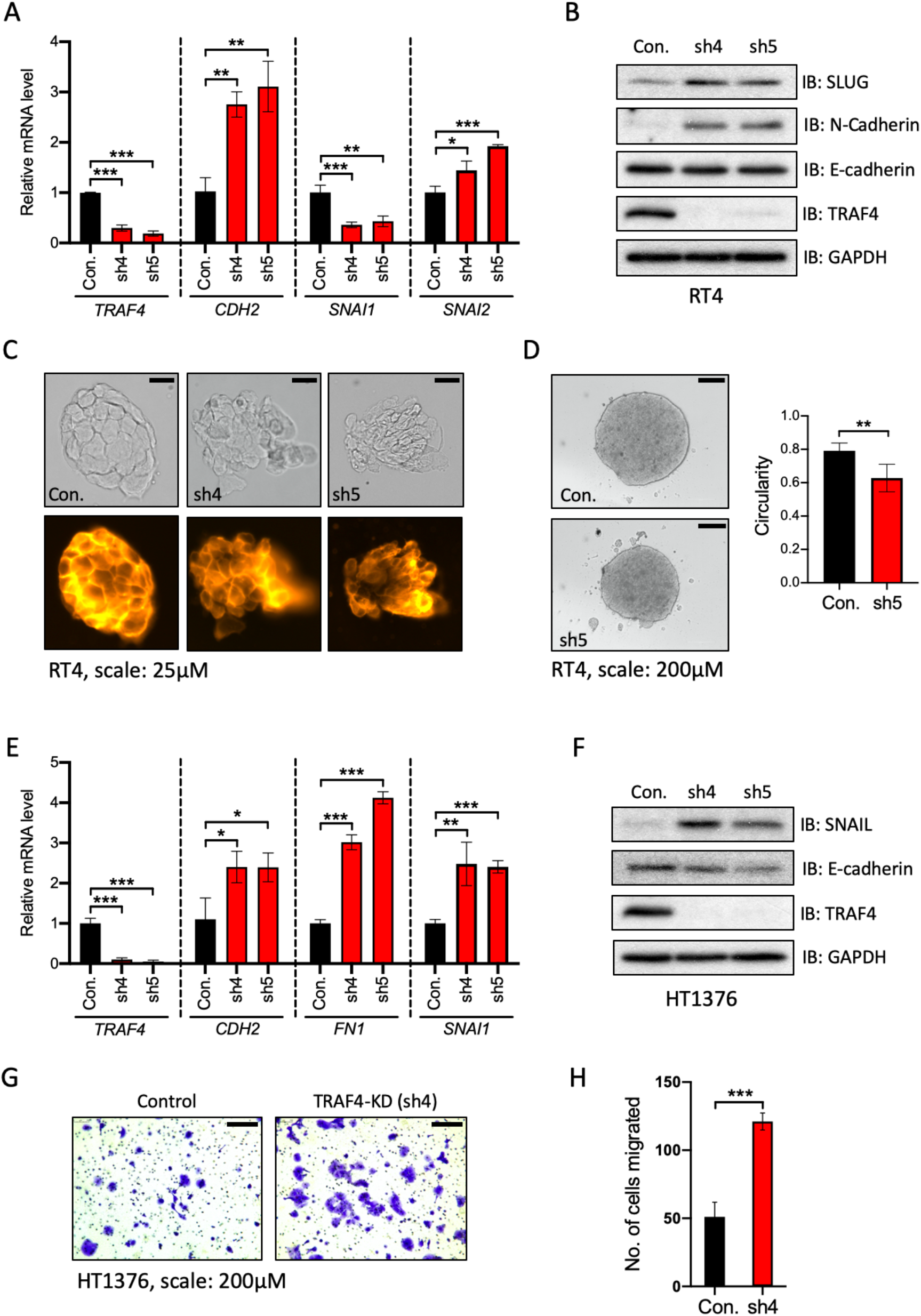
Knockdown of TRAF4 in epithelial (bladder cancer) cell lines leads to loss of epithelial integrity and changes in EMT marker expression. **A** Real-time PCR result from RT4 cells showing mRNA expression levels of genes indicated; error bars represent ± SD, \**P* ≤ 0.05, \*\**P* ≤ 0.01, \*\*\**P* ≤ 0.001 calculated using two-tailed student’s t test. **B** Immunoblot result showing EMT markers protein expression changes in RT4 cells upon TRAF4 knockdown. **C** RT4 cell colonies imaged by brightfield (top panels) or after exposure to CellMask™ Orange plasma membrane stain (bottom panels), scale: 25μM. **D** Images depicting RT4 spheroids made from control (empty vector pLKO) and TRAF4 knockdown cells (sh5), scale: 200μM. The graph represents circularities calculated from five independent spheroids of different sizes. Error bars represent ± SD, \*\**P* ≤ 0.01 calculated using two-tailed student’s t test. **E** Real-time PCR result from HT1376 cells showing mRNA expression of the genes indicated; error bars represent ± SD, \**P* ≤ 0.05, \**P* ≤ 0.05, \*\**P* ≤ 0.01, \*\*\**P* ≤ 0.001 calculated using two-tailed student’s t test. **F** Immunoblot result showing EMT markers protein expression changes in HT1376 cells upon TRAF4 knockdown. **G** Representative images of transwell assays performed on HT1376 cells, cells were stained with crystal violet, scale: 200μM. **H** Quantification of number of migrated cells from four random fields; error bars represent ± SD, \*\*\**P* ≤ 0.001 calculated using two-tailed student’s t test.

To further examine if TRAF4 knockdown affects the 3-dimensional architecture of cell structures, RT4 cell spheroids were generated. Spheroids recapitulate tumour cell clusters and can be considered in many ways a better model representative of an *in vivo* situation than 2-dimensional cell culturing. Fig 3D depicts cell spheroids made from control RT4 (Empty vector pLKO) and TRAF4 knockdown cells. We demonstrate here that in the TRAF4 knockdown spheroids, several clumps of cells were dissociated or excluded from the main spheroids bodies. Moreover, spheroids in the knockdown group were more irregular in shape compared to control as measured through its circularities. Such cell-exclusion phenotype has been observed in previous studies, which are reflective of loss of certain tight junction components in epithelial cells[20, 21]. We further demonstrate the effects of TRAF4 knockdown using yet another epithelial cell line, HT1376. TRAF4 knockdown using 2 different shRNAs (Sup. Fig 3C) led to an increase in mRNA expression levels of mesenchymal markers *CDH2*, *FN1* (encoding Fibronectin) and *SNAI1* (Fig 3E). Although we saw a concomitant increase in the EMT-TF SNAIL (Fig 3F), we couldn’t observe N-cadherin levels through immunoblotting as its levels were too low to be detected. Moreover, levels of E-cadherin remained at best unchanged upon TRAF4 knockdown in these cells. Previous studies have demonstrated invasive properties of these cells[22]. We hypothesized that since EMT has been linked to migratory and invasive properties, knockdown of TRAF4 would lead to enhanced invasive behaviour in these cells. As can be seen in Fig 3G and 3H, knockdown of TRAF4 indeed increased the number of migrated cells as determined by transwell assays. Based on our observations and results so far, we are able to conclude that knock down of TRAF4 in epithelial cells indeed disrupts its epithelial-architecture and organization. Importantly, it leads to an increase in expression of major EMT-TFs such as SNAIL and SLUG. When we compared *SNAI1* and *SNAI2* expression across 5 bladder cancer cells lines, we observed an almost mutually exclusive expression pattern of these transcription factors (Sup. Fig 3D and 3E), suggesting a certain level of functional redundancy or compensation in the bladder cancer cell lines that were tested.

### TRAF4 targets SMURF1 for ubiquitination and degradation

We hypothesized that TRAF4 is destabilizing SMURF1 levels and that upon TRAF4 knockdown, levels of SMURF1 increase. Of note, SMURF1 has been previously shown to be a positive regulator of EMT progression[23]. Fig 4A shows that in RT4 cells, upon TRAF4 knockdown, levels of SMURF1 indeed increase compared to control cells. This is also observed in HT1376 (Fig 4B). Importantly, SMURF1 mRNA levels do not change upon TRAF4 knockdown in both these cell lines (Fig 4C), suggesting that TRAF4 could be targeting SMURF1 protein for degradation. We further supported this notion by over-expressing SMURF1 along with TRAF4 or its inactive mutants in HEK293T to investigate the interplay. As can be seen in Fig 4D, the combined expression of TRAF4 and SMURF1 leads to decreased levels of SMURF1. Importantly a catalytically inactive version of TRAF4 (C/A) or the RING deletion mutant did not achieve the same effect. Interestingly, ectopic expression of TRAF4, but not it’s catalytically inactive mutant (C/A) leads to enhanced poly-ubiquitination of SMURF1 (Fig 4E). Taken together, our results suggest that TRAF4 is capable of targeting SMURF1 for ubiquitination mediated degradation. The role of SMURF1 in cancer cell invasion is well-documented[24]. To understand if SMURF1 plays a more notorious role in mesenchymal (bladder cancer) cells due to reduced expression of TRAF4, we utilized the MBT-2 cell line for our investigation. The MBT-2 cell line is an established mouse bladder cancer cell line with highly invasive properties. As can be seen in Sup Fig 4A, levels of Traf4 are quite low in MBT-2 cells compared to other cell lines in our investigation. Also, levels of mesenchymal markers (N-cadherin and Vimentin) appeared to be highly expressed while epithelial markers (E-cadherin) is expressed poorly in this cell line. To test the effects of Smurf1 during invasion of these cells, we stably knockdown Smurf1 using two independent shRNAs (Fig 4F), which showed the best knockdown efficiency (Sup. Fig 4B). We observed that upon Smurf1 knockdown, there was significantly lower number of migrated cells compared to vector control (Fig 4G and 4H). We tested the effects of SMURF1 inhibition through a commercially available SMURF1 inhibitor, A01. As can be seen in Fig 4I and 4J, treating T24 cells with A01 significantly reduced the wound healing ability and rate of migration compared to control cells (DMSO treated). Of note, this SMURF1 inhibitor does not target the catalytic activity, but targets the ability of SMURF1 to bind to and subsequently degrade SMAD1 and SMAD5, which are intracellular effectors of BMP[25]. Also, MTS cell viability assay revealed that the inhibitor had slightly enhanced the proliferative capacity of these cells (Fig 4K). Taken together, our results demonstrate that TRAF4 can indeed inhibit SMURF1 protein levels in bladder cancer cells and that SMURF1 enhances migration and invasion of these mesenchymal bladder cancer cells.

**Figure 4.**
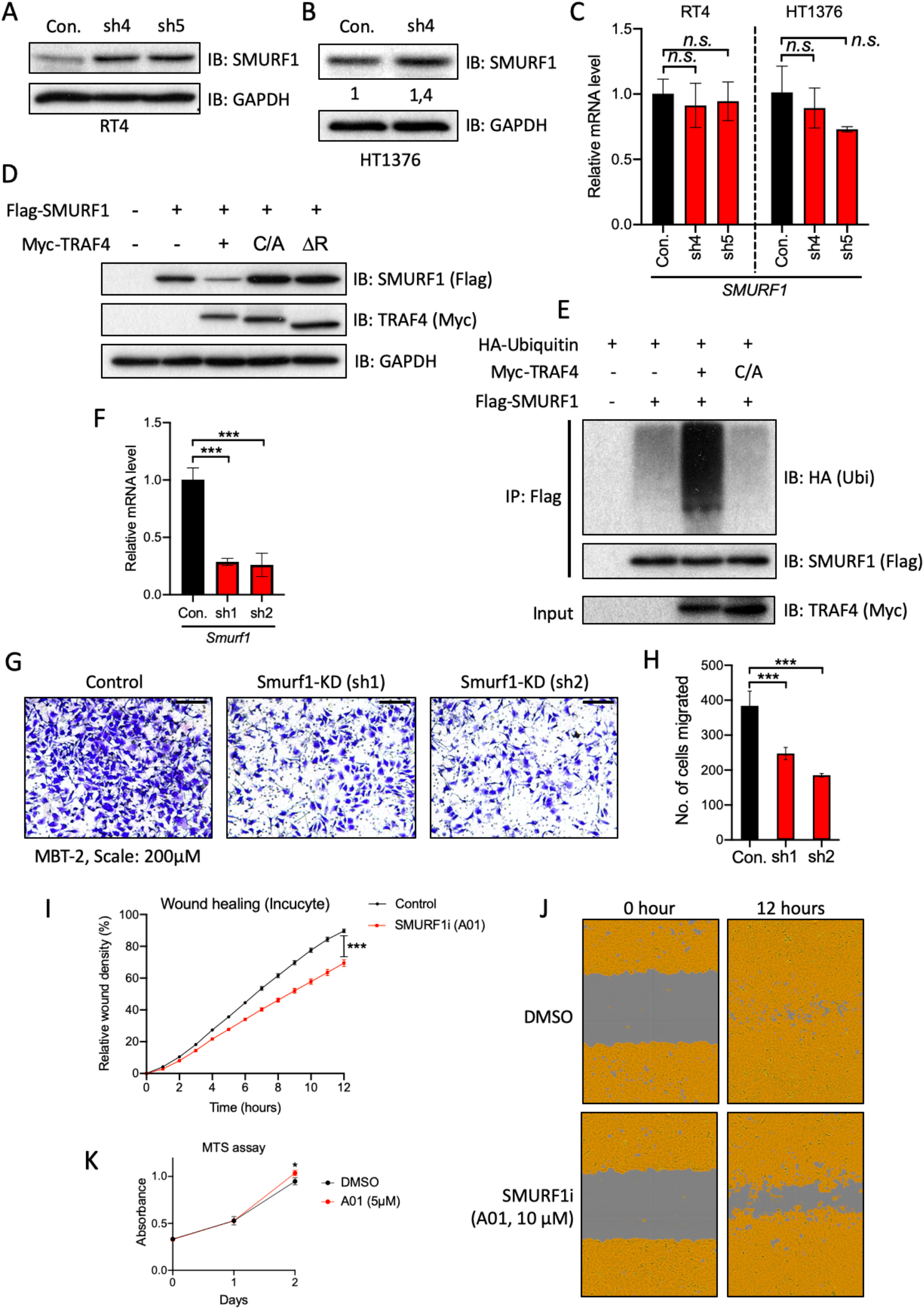
TRAF4 targets SMURF1 for ubiquitination and degradation. **A** Immunoblot result of RT4 control (empty vector pLKO) and TRAF4 knockdown cells (sh4 and sh5) probed with the indicated antibodies. **B** Immunoblot result of HT1376 control (empty vector pLKO) and TRAF4 knockdown cells (sh4) probed with the indicated antibodies. Numbers represent relative quantification of SMURF1 levels with respect to GADPH. **C** Real-time PCR result showing *SMURF1* mRNA expression levels in RT4 and HT1376 (control and TRAF4 knockdown) cells; error bars represent ± SD, *n.s.* indicates non-significant P value. **D** Immunoblot result from 293T cells transfected with the indicated plasmids. **E** Ubiquitination assay performed in 293T cells with Flag antibodies, over-expressing the indicated plasmids. Cells were treated with MG132 (2μM) overnight prior to lysis. Representative result from three independent experiments. **F** Real-time PCR showing *Smurf1* mRNA levels in MBT-2 control (pLKO vector) and Smurf1 knockdown cells (sh1 and sh2); error bars represent ± SD, \*\*\**P* ≤ 0.001 calculated using two-tailed student’s t test. **G** Representative images of transwell assay performed on MBT-2 control and Smurf1 knockdown cells, which were stained with crystal violet, scale: 200μM. **H** Quantification of number of migrated cells from four random fields, error bars represent ± SD, \*\*\**P* ≤ 0.001 calculated using two-tailed student’s t test. **I** Graph showing relative wound density from Incucyte, images were obtained every 1 hour after wound was produced, T24 cells were treated with either DMSO or SMURF1 inhibitor, A01 (10μM); error bars represent ± SEM, \*\*\**P* ≤ 0.001 calculated using two-tailed student’s t test. Representative result from three independent experiments**. J** Representative images from graph in I, brown area represents the cell coverage and grey area is the wound produced and remaining. **K** MTS assay performed on T24, cells were treated with either control (DMSO) or SMURF1i, A01 at 5μM. Absorbance was read at the indicated time points; error bars represent ± SD from three sample replicates, \**P* ≤ 0.05 calculated using two-tailed student’s t test.

### Stable over-expression of TRAF4 diminishes the migratory and invasive properties of mesenchymal cells

We next assessed if ectopic expression of TRAF4 in mesenchymal cell lines affect their functional properties. To that end, we stably over-expressed either empty vector (myc-tag), TRAF4 or TRAF4 (C/A) as shown in Fig 5A. It’s interesting to note here that the SMURF1 levels were down-regulated in TRAF4 expressing cells but not in TRAF4 (C/A) cells. From Fig 5B and 5C, we demonstrate that TRAF4 cells have reduced number of migrated cells. Also, the wound-healing ability and rate of migration was affected in TRAF4 cells compared to its vector control (Fig 5D and 5E). Interestingly, however, TRAF4 expressing cells had enhanced proliferative growth capacities (Fig 5F). We extended our studies to MBT-2 mesenchymal cell line. Similarly, we stably over-expressed TRAF4 in these cells (Fig 5G). Ectopic expression of TRAF4 in MBT-2 cells diminished their invasive abilities (Fig 5H and 5I). On closer examination, morphologically, TRAF4 over-expressing cells tended to cluster together than its control cells, especially in low seeded culture conditions (Sup. Fig 4C). Collectively, we demonstrate that TRAF4 over-expression dampens some of the aggressiveness of mesenchymal cell lines.

**Figure 5.**
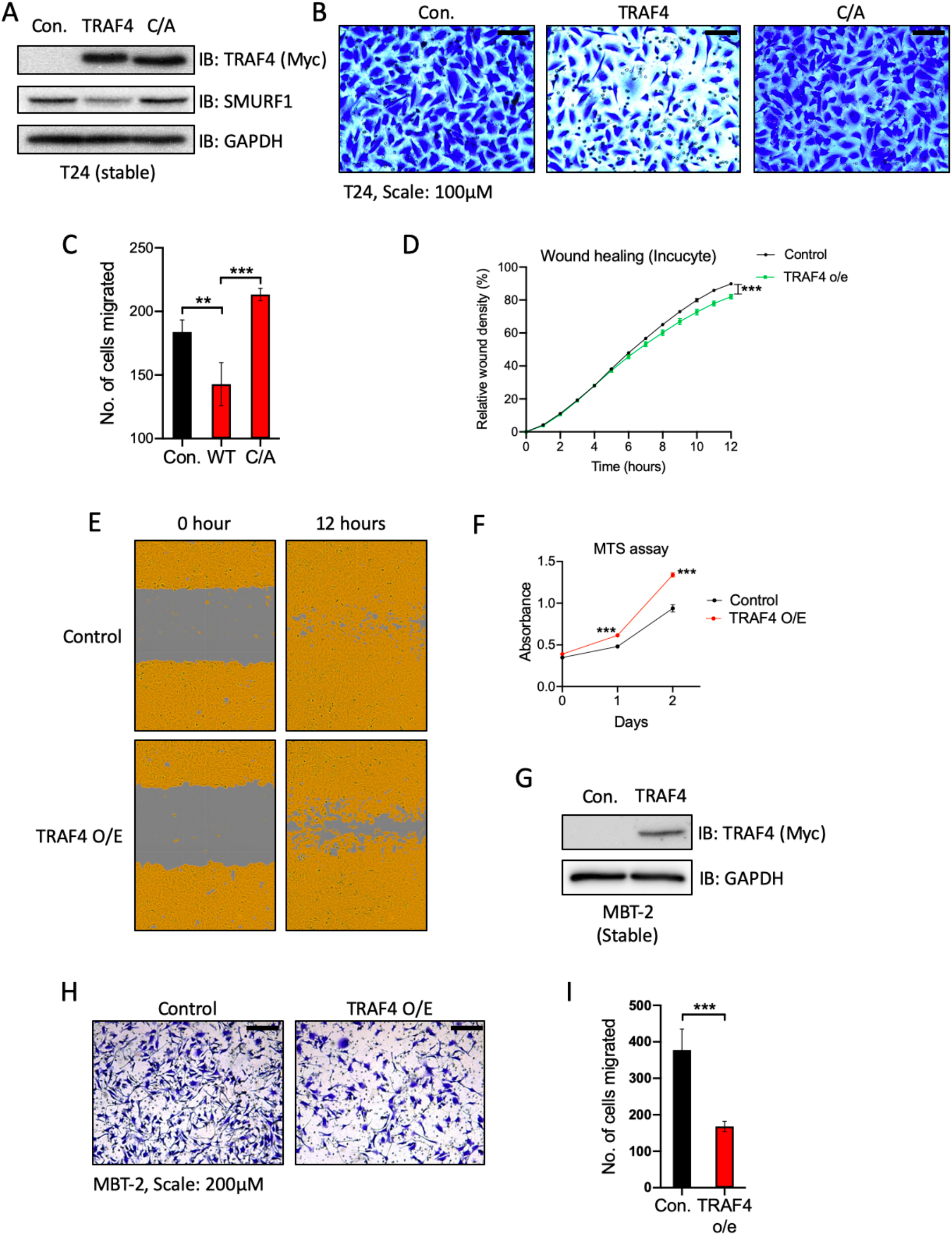
Reintroducing TRAF4 in mesenchymal cells negatively affects their migratory and invasive properties. **A** Immunoblot showing T24 cells stably expressing either control vector (myc-tag), TRAF4 or catalytically inactive TRAF4 mutant (C/A: cysteine to alanine at residue C18). **B** Representative images of transwell assays performed on T24 cells, cells were stained with crystal violet, scale: 100μM. **C** Quantification of number of migrated cells from four random fields, error bars represent ± SD, \*\**P* ≤ 0.01, \*\*\**P* ≤ 0.001 calculated using two-tailed student’s t test. **D** Graph showing relative wound density result from Incucyte. Representative result from three independent experiments; error bars represent ± SEM, \*\*\**P* ≤ 0.001 calculated using two-tailed student’s t test. **E** Representative images from graph in D, brown area represents the cell coverage and grey area is the wound produced and remaining. **F** MTS assay performed with either control or TRAF4 stably expressing T24 cells. Absorbance was read at the indicated time points; error bars represent ± SD from three sample replicates, \*\*\**P* ≤ 0.001 calculated using two-tailed student’s t test. **G** Immunoblot result showing MBT-2 cells stably expressing either control (empty vector with Myc-tag) or Myc-TRAF4. **H** Representative images of transwell assays performed on MBT-2 control and TRAF4 over-expressing cells, which were stained with crystal violet, scale: 200μM. **I** Quantification of number of migrated cells from four random fields, error bars represent ± SD, \*\*\**P* ≤ 0.001 calculated using two-tailed student’s t test.

### Mis-expression of TRAF4 in bladder cancer cell lines affects NF-κB and BMP signalling pathways

We then sought to determine if there are some central or common TRAF4 regulated signalling pathways in bladder cancer cells. To that end, we stably over-expressed TRAF4 in T24 cells (Sup. Fig 4D), performed transcriptomic (RNA-seq) analysis and looked for changes in gene signatures in the eleven most commonly studied cancer causing signalling pathways as shown. Nine out of the eleven signalling pathways showed some changes in enrichment scores to varying degrees (Sup. Fig 4E). The most affected was the NF-κB signalling pathway exhibiting a strong decrease in activity, followed by the BMP/SMAD signalling pathway with an increase (Fig 6A). We also looked at changes in gene expression when we knocked down TRAF4 in HT1376 using two independent shRNA hairpins. As shown in Fig 6B, there were 252 genes that were upregulated upon TRAF4 knockdown in HT1376 (common in both shRNA hairpins) and reciprocally downregulated in T24 upon TRAF4 over-expression (Sup. Table 3). Similarly, we detected 96 genes that were up-regulated in T24 and down-regulated in HT1376 compared to their respective controls (Fig 6C and Sup. Table 3). We observed five genes from the BMP gene signature that were reciprocally regulated between the two cell lines, *ID1, ID2, ID3, DKK1* and *TNFRSF11B* (Fig 6D). It’s interesting to note that *ID1, ID2* and *ID3,* which are bona fide genetic targets of BMP signalling[26], get down-regulated upon TRAF4 knockdown in HT1376 cells and up-regulated upon TRAF4 over-expression in T24 cells. Upon examining an NF-κB gene signature within the reciprocally regulated genes, we found four common genes as shown in the heatmap (Fig 6D). Importantly, the majority of NF-κB genes were down-regulated upon TRAF4 over-expression in T24 with very few genes being reciprocally up-regulated in HT1376. This suggests that unlike the BMP signalling pathway, NF-κB has a more prominent role to play in mesenchymal cells. Since our observations regarding TRAF4 strongly correlated with the EMT status of cells, we also investigated the EMT gene signature. Upon closer examination of the EMT gene signature within the reciprocally regulated genes between the two cell lines, we found four genes as shown (Fig 6D). Interestingly, *FN1* (encoding Fibronectin) and *ITGAV* (encoding Integrin subunit α-V) have been well-documented as potential biomarkers or targets for bladder carcinoma[27, 28].

**Figure 6.**
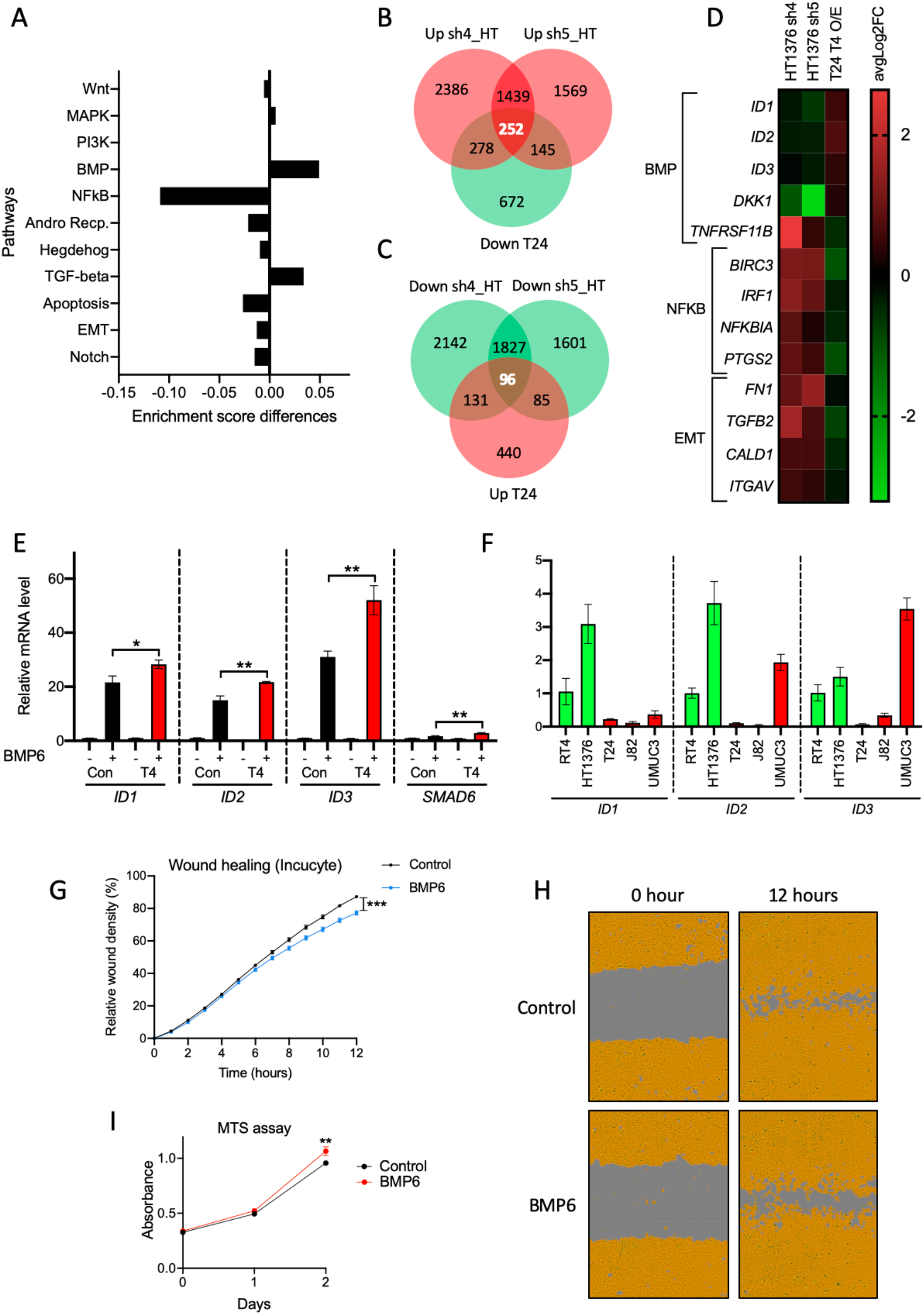
Mis-expression of TRAF4 in bladder cancer cell lines affects downstream BMP signalling target genes. **A** Graph showing differences in enrichment scores when TRAF4 is over-expressed in T24 cells. Gene signatures of eleven major cancer associated signalling pathways were considered for analysis. **B** Venn diagram showing number of genes which are up-regulated in two independent hairpins targeting TRAF4 in HT1376 (pink) and down-regulated with stable over-expression of TRAF4 in T24 (green) compared to their respective controls. 252 genes in the middle represents the reciprocally affected common genes. Results were obtained from four independent replicates for each sample. **C** Venn diagram showing number of genes which are down-regulated in two independent hairpins targeting TRAF4 in HT1376 (green) and up-regulated with stable over-expression of TRAF4 in T24 (pink) compared to their respective controls. 96 genes in the middle represents the reciprocally affected common genes. Results were obtained from four independent replicates for each sample. **D** Heatmap showing the common mis-regulated genes within the BMP, NF-κB and EMT gene signatures in both cell lines. **E** Real-time PCR result showing mRNA expression levels of the indicated genes in control (empty vector myc-tag) vs TRAF4 over-expressing T24 cells upon stimulation of BMP6 (50ng/ml) for 1 hour, error bars represent ± SD, \**P* ≤ 0.05, \*\**P* ≤ 0.01, calculated using two-tailed student’s t test. **F** Real-time PCR showing mRNA expression levels of *ID1, ID2* and *ID3* in the indicated cell lines. **G** Graph showing relative wound density result from Incucyte, images were obtained every 1 hour after wound was produced, T24 cells were treated with BMP6 (50ng/ml); error bars represent ± SEM, \*\*\**P* ≤ 0.001 calculated using two-tailed student’s t test. **H** Representative images from graph in G, brown area represents the cell coverage and grey area is the wound produced and remaining. **I** MTS assay performed with T24 cells, either control and stimulated with BMP6 (50ng/ml). Absorbance was read at the indicated time points; error bars represent ± SD from three sample replicates, \*\**P* ≤ 0.01 calculated using two-tailed student’s t test.

Next, we sought to determine the consequences of BMP stimulation on mesenchymal cells. Interestingly, we observed that the canonical BMP pathway target genes (*ID1, ID2, ID3* and *SMAD6* [26, 29]) are expressed at higher levels upon BMP6 stimulation in T24 cells stably expressing TRAF4 (Fig 6E). This effect was also observed in MBT-2 upon BMP6 stimulation (Sup. Fig 5A). Next, a BRE-luciferase reporter assay was used to measure downstream BMP signalling activity in 293T cells[30]. As can be seen from Sup. Fig 5B, BMP6 stimulation led to significantly higher luciferase activity in cells transfected with TRAF4. We then hypothesized that perhaps these three *ID* genes could be expressed at a higher level in epithelial cell lines due to differential TRAF4 expression, and we observed that indeed in RT4 and HT1376 cell lines, their levels were generally higher than in mesenchymal cell lines (Fig 6F). The wound-healing ability and rate of migration was reduced in BMP6 stimulated cells compared to control (Fig 6G and 6H). Interestingly again, BMP6 stimulated cells slightly increased the proliferative growth capacities (Fig 6I). Moreover, we also observed that the BMP receptor kinase inhibitor (LDN193189) rescued the inhibitory effects of TRAF4 on the wound-healing ability (Sup. Fig 5C and 5D). Taken together, our results shed light on the potential role of BMP signalling in aggressive bladder cancer cells.

### TRAF4 expression positively correlates with pSMAD1/5/8 levels and negatively with p-p65 levels

Since we previously observed that TRAF4 can negatively influence the NF-κB gene signature and positively affect BMP/SMAD genes, we hypothesized that perhaps the NF-κB signalling pathway can also have a negative influence on the BMP/SMAD pathway and TRAF4 can rescue this effect. To that end, we evaluated this relationship using a BRE-luciferase reporter activity. As can be seen in Fig 7A, combined stimulation of BMP and TNF-α negatively affects the signalling output. Interestingly, over-expression of TRAF4 rescued the negative effect of TNF-α on the BMP pathway. Similar effects were observed when T24 cells were stimulated with a combination of BMP and TNF-α and *ID* genes expression were analysed (Fig 7B). Consistent with previous observations, over-expression of TRAF4 negatively affected the NF-κB reporter activity (Sup. Fig 5E) [31]. To further extend our observations and if it can be translated into analysis of patient material, we performed immunohistochemistry on tissue microarray samples obtained from Biomax U.S (BL802b). For the BMP signalling pathway, validated pSMAD1/5/8 antibodies and for NF-κB, validated p-p65 antibodies were used as readout. We observed a significant positive correlation between TRAF4 and pSMAD1/5/8 levels (Fig 7C and 7E). And interestingly, a significant negative correlation between TRAF4 and p-p65 levels (Fig 7D and 7E). Taken together, our findings confirmed that repression of TRAF4 enhances NF-κB activity and decreases BMP activity in bladder cancer.

**Figure 7.**
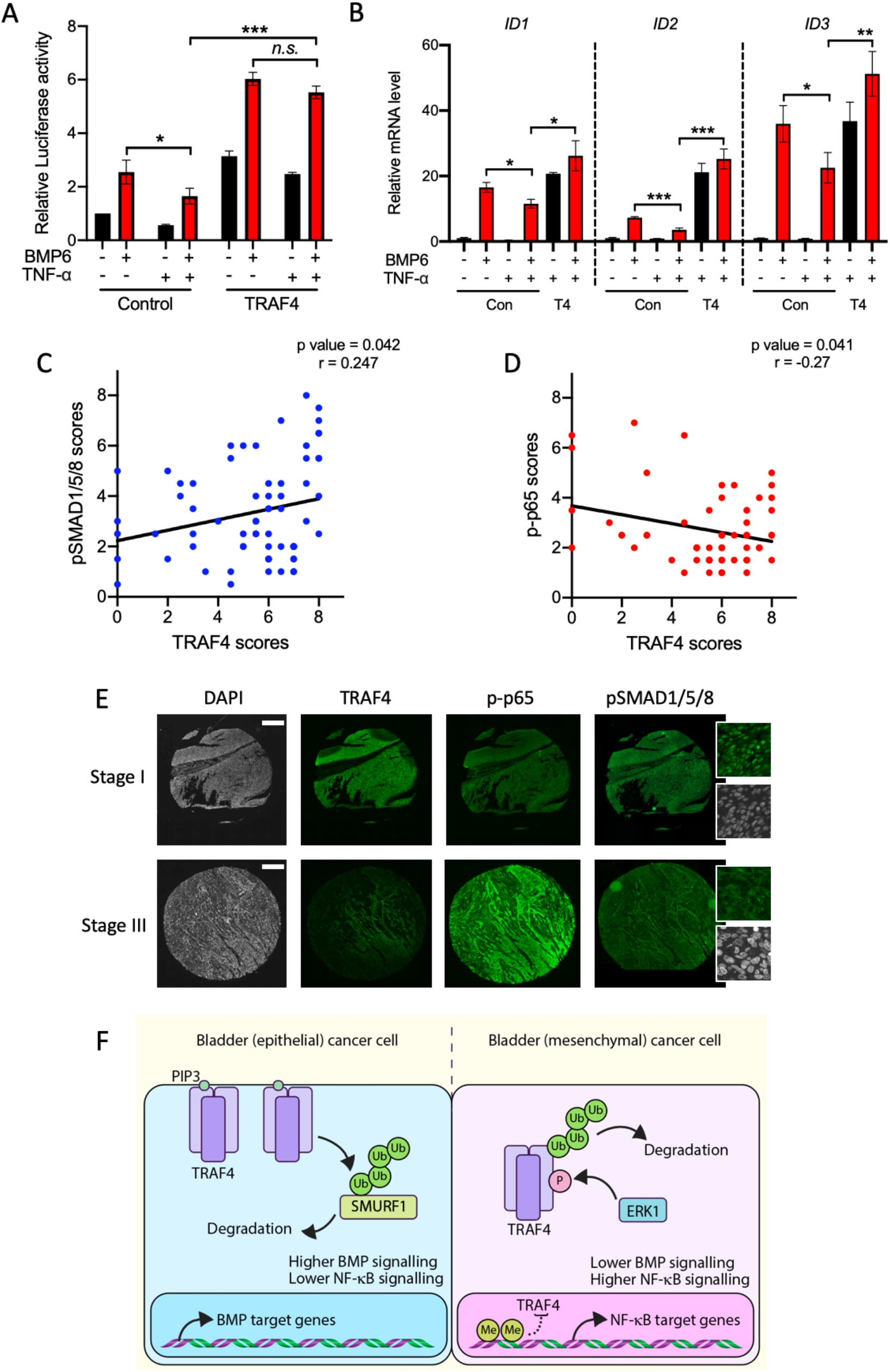
TRAF4 expression correlates positively with pSMAD1/5/8 levels and negatively with p-p65 levels in bladder tumours. **A** Luciferase reporter assay in 293T transfected with BRE-luciferase reporter, SV40 Renilla and either empty vector control or TRAF4, transfected cells were stimulated overnight with BMP6 (50ng/ml) and/or TNF-α (10ng/ml) where indicated, error bars represent ± SD, \**P* ≤ 0.05, \*\*\**P* ≤ 0.001, calculated using two-tailed student’s t test, *n.s.* indicates non-significant P value. Representative result from three independent experiments. **B** Real-time PCR result showing mRNA expression levels of the indicated genes in control (empty vector myc-tag) vs TRAF4 over-expressing T24 cells upon stimulation of BMP6 (50ng/ml) and/or TNF-α (10ng/ml) where indicated for 1 hour. Error bars represent ± SD, \**P* ≤ 0.05, \*\**P* ≤ 0.01, \*\*\**P* ≤ 0.001, calculated using two-tailed student’s t test. **C** Regression analysis showing correlation between TRAF4 expression levels (scores) and pSMAD1/5/8 scores from bladder cancer patients. Pearson’s test was used to determine the correlation between TRAF4 and pSMAD1/5/8 scores. **D** Regression analysis showing correlation between TRAF4 expression levels (scores) and p-p65 scores from bladder cancer patients. Pearson’s test was used to determine the correlation between TRAF4 and p-p65 scores. **E** Representative images of continuous sections of tissue microarray samples probed with the indicated antibodies using fluorescent immunohistochemistry. The magnified insets for pSMAD1/5/8 shows nuclear staining. **F** Schematic representation of TRAF4 signalling dynamics between epithelial and mesenchymal bladder cancer cells. Ub denotes Ubiquitin, Me is methylated promoter DNA and P stands for Phosphorylation of S334 site.

## Discussion

The relationship between TRAF4 and cancer progression has been well documented[32]. In our study, we investigated the role of TRAF4 during bladder cancer progression and observed strong correlations between its expression and overall patient survival. Data mining from publicly available patient material and immunohistochemistry results revealed that TRAF4 expression gradually gets lowered as the cancer progresses. Pathologically, stage 2 and 3 bladder tumours are more advanced and have muscle invasiveness compared to stage 1 tumours and consistently, we observed higher TRAF4 levels in stage 1 tumours. Furthermore, in both cultured cells and bladder cancer patient material, we observed strong links of TRAF4 expression with BMP/SMAD and NF-κB signalling pathways. Mechanistically, we show that TRAF4 ubiquitinates and degrades SMURF1, which is considered to be a pro-EMT and oncogenic protein. Our results further gives credence to the claim that TRAF4 enhances BMP/SMAD signalling as SMURF1 is a negative regulator (Fig 7F).

Epithelial-Mesenchymal Transition is a prominent event during cancer metastasis, especially in the case of bladder carcinoma where (epithelial) cancer cells usually have to gain mesenchymal properties to make their way through the bladder muscle wall. Interestingly, we found a strong correlation between the EMT status of bladder cancer cells and TRAF4 expression. Importantly, knock down TRAF4 in an epithelial cell line, RT4 led to loosely attached cells from colonies suggesting possible loss of epithelial tight junction components. In our EMT score analysis, the RT4 cell line apparently had the lowest score which gave it the highest epithelial status. Interestingly, depletion of TRAF4 in a commonly used epithelial breast cancer cell line, MCF10 did not show similar results[16]. We also detected higher *TRAF4* levels in MDA-MB-231, a breast cancer (mesenchymal) cell line compared to MCF10A[6]. Our observations have uncovered a deeper understanding about epithelial behaviour of bladder cells from other cancer types. Physiologically, bladder urothelial cells have high expressions of TRAF4 that perhaps enables the bladder to maintain a strong barrier against stored urine. Changes in expression of EMT transcriptions factors, SNAIL and SLUG were observed upon TRAF4 knockdown, for example in RT4, an increase in SLUG and a decrease in SNAIL and *vice versa* in HT1376. It was interesting to see such opposite trends on the two transcription factors using the two epithelial cell lines, although such reciprocal effects of SNAIL and SLUG expression have been previously documented[33].

We initially hypothesized that *TRAF4* is epigenetically repressed due to lower mRNA levels in the more aggressive cells. Epigenetic repression of genes is commonly seen during cancer progression and is carried out by DNA methylation enzymes that methylate certain regions on promoters to diminish their transcriptional activities[34]. Interestingly, our results also show that TRAF4 is unstable at protein levels in more aggressive bladder cancer cells. We found an ERK kinase phosphorylation site that not only is responsible for TRAF4 protein stability but also its localization to plasma membrane. TRAF4 molecules form a trimer and bind to PIP3 at the cell surface to maintain tight junction stability; phosphorylation at a serine residue (S334) close to PIP3 binding sites creates an overall negative environment that repels it from the plasma membrane. Perhaps this doesn’t come as a surprise as many of these aggressive (bladder) cancer cells have mutations in components of MAPK pathways such as Raf or Ras that increases activity of downstream ERK signalling.

In our investigation, we observed that TRAF4 affected the endogenous (protein) levels of SMURF1 in bladder cancer cells, TRAF4 keeps SMURF1 levels in check and as TRAF4 levels get repressed, SMURF1 becomes more active and stable. We have previously observed that TRAF4 is able to ubiquitinate SMURF2, thereby potentiating TGF-β signalling that enhances breast cancer metastasis[6]. SMURF1 has high sequence similarity to SMURF2 and also belongs to the same E3 ubiquitin ligase sub-family (HECT-domain, NEDD4 sub-group). Additionally, there have been other important studies reflecting the dynamic interplay between TRAF4 and SMURF1[18, 35]. Our observations reveal that up-regulation of SMURF1 due to reduced TRAF4 expression in later stages of bladder cancer progression could potentially dampen the BMP signalling output.

We performed transcriptomic and pathway analysis in an unbiased manner, which revealed TRAF4 affected the NF-κB and BMP pathways in bladder cell lines. The NF-κB pathway has been previously shown to be negatively affected by TRAF4[36], and has been documented to suppress apoptosis and enhance cell-proliferation during bladder cancer progression[31]. The pathway also has strong ties with immune response and inflammation. The BMP pathway on the other hand is positively associated with TRAF4 expression in cells and patient material. Our novel observations further revealed a strong link to BMP signalling target genes in particular *ID1, ID2* and *ID3,* which are bona fide downstream targets. These results are consistent with the notion that SMURF1 negatively regulates BMP signalling components i.e. SMAD1/5 and BMP receptors[37–39]. Interestingly, the NF-κB pathway in itself could also dampen the BMP signalling output, an effect that can be blocked by TRAF4. Addition of BMP ligand to mesenchymal cells had an inhibitory effect on migration and a SMURF1 inhibitor (A01) could also give similar results. Apparently, the compound can inhibit SMURF1 from degrading SMAD1 and SMAD5 components of BMP signalling pathway rather than its catalytic activity.

In summary, our investigation uncovers a negative role for TRAF4 in bladder cancer progression, in contrast to other cancers that were examined. Furthermore, we demonstrate the absence of TRAF4 expression as a biomarker to detect aggressive types of bladder cancers. Moreover, even though increase in TRAF4 expression is seen in a lot of cancers, caution must be taken in the potential development or use of a TRAF4 inhibitor in future. Furthermore, coupling SMURF1 inhibitor (A01) or use of BMP agonists in bladder cancers with low TRAF4 expression will be interesting to explore.

## Materials and methods

### Cell culture conditions

Bladder cancer cell lines and HEK293T were purchased from ATCC (American Type Culture Collection). Cells were grown in Dulbecco’s modified eagle medium (DMEM) supplemented with 10% fetal bovine serum and penicillin/streptomycin. Cells were regularly tested for the absence of mycoplasma contamination and were genotyped and authenticated. Cells were grown in 5% CO_2_ atmosphere incubator at 37°C. Where appropriate, cells were treated with BMP6 (50ng/ml), TNF-α (10ng/ml) MG132 (2μM), LDN193189 (120nM), SMURF1i-A01 (5 or 10μM), MEK (PD0325901, 2μM), 5’-Azacytidine (5μM) and Cycloheximide (10μg/ml) for the indicated hours.

### Transient transfection

HEK293T cells were transfected with the indicated plasmids using calcium chloride and HEPES buffered saline (pH 6.95). After an overnight incubation, cells were washed twice with 1X PBS solution and replenished with fresh serum containing media. HT1376 cells were transfected using lipofectamine 2000 according to manufacturer’s protocol. The following vectors and its derivates were used: pcDNA3.1 (6xMyc) TRAF4, pFLAG-CMV SMURF1, GFP-ERK1 was a gift from Rony Seger (Addgene plasmid # 14747)[40] and GFP-TRAF4 was a gift from Ying Zhang (Addgene plasmid # 58318)[18].

### Stable transfection

Stable knockdown of TRAF4 in RT4, HT1376 or over-expression in T24 cells were generated using short hairpin RNAs through lentiviral transduction. HEK293T cells were transfected with pLKO.1 puro vectors (Sigma Mission shRNAs) or pLV-IRES Lenti Puro TRAF4 along with lentiviral packaging plasmids (pCMV-VSVG, pMDLg-RRE and pRSV-REV). The media containing viral particles were collected 48 hours later and passed through 0.45μM filter. The supernatant with Polybrene (0.01%) was used to transduce bladder cancer cells. The cells were further selected with Puromycin (1μg/ml) containing medium. The list of short hairpins used for knockdown are provided in Sup. Table 6.

### Immunofluorescence

Labelling of plasma membrane was achieved with CellMask™ Orange plasma membrane stain (ThermoFisher) solution. RT4 cells were treated with the solution according to manufacturer’s instructions and images were captured soon after using Leica fluorescence microscope. HT1376 transfected with expression constructs for GFP, GFP-TRAF4 or its mutants, and transfected cells were visualized under fluorescence microscope and images were captured. Presence of GFP signal at the membrane was quantified using ImageJ analysis.

### *In vivo* phosphorylation experiment

Transfected HEK293T cells were lysed in ELB buffer (250mM NaCl, 0.5%Nonidet P-40, HEPES 50mM, pH 7.3), supplemented with protease inhibitors and serine/threonine phosphatase inhibitors: 50mM sodium flouride and 10mM β-glycerophosphate. Protein concentration was estimated on the lysates using a bicinchoninic acid protein assay Kit (5000111, Bio-Rad). Equal protein concentrations for each of the samples were incubated with Myc antibodies overnight, followed by incubation with Protein G beads for 1 hour at 4°C. After several washing steps with 1XPBS, beads were boiled in 2X sample buffer. The resulting supernatants were processed for immunoblotting.

### Culturing of cell spheroids

Cell spheroids were generated using RT4 cells. 1.5% agarose was boiled until it dissolved in 1XPBS, then added onto a sterile 96-well plate (100μl each well) and it was let to solidify. About an hour later, RT4 cells were trypsinized, counted and diluted in media. About 200μl of media containing the appropriate amount of cells were added onto the agarose beds formed on the 96-well plate. The plate was spun down at 1000 RPM for 2 minutes and incubated at 37°C CO_2_ incubator overnight. The following day, cell spheroids were checked under a Leica microscope. Spheroids were assessed for circularity using ImageJ software.

### *In vivo* ubiquitination assay

Cells were washed in ice cold 1XPBS (twice) and then lysed with RIPA buffer (25mM Tris HCl, pH 7.4, 150mM NaCl, 1%Nonidet P-40, 1%SDS, 0.5%sodium deoxycholate), supplemented with protease inhibitors and 10mM N-ethylmaleimide. Lysates were sonicated, boiled at 95°C for 5 minutes and diluted with RIPA buffer containing 0.1% SDS. Lysates were centrifuged at 4°C for 15 minutes. Thereafter, protein estimation was performed and equal amounts of lysates were incubated with Myc antibodies overnight, followed by incubation with Protein G beads for 1 hour at 4°C. After several washing steps with 1XPBS, beads were boiled in 2Xsample buffer. The resulting supernatant was processed for immunoblotting.

### Transcriptomics, gene signatures, pathway analysis and enrichment scores

TRAF4 was knocked down using shRNA in HT1376 using lentiviral transduction. Cells with empty vector (pLKO) was used as control. Four independent experimental replicates were used for each condition. T24 cells stably over-expressing Myc-TRAF4 or empty vector (Myc-tag) were generated. Again, four independent experimental replicates were used. The cells were processed for RNA extraction and sent to BGI Tech (Hong Kong) for further processing. RNA-Seq files were processed using the opensource BIOWDL RNAseq pipeline version 3.0.0 (https://zenodo.org/record/3713261#.X4GpD2MzYck) developed at the LUMC. The pipeline performs FASTQ pre-processing (including quality control, quality trimming, and adapter clipping), RNA-seq alignment, read quantification, and optionally transcript assembly. FastQC was used for checking raw read QC. Adapter clipping was performed using Cutadapt (v2.8) with default settings. RNA-Seq reads’ alignment was performed using STAR (v2.7.3a) on GRCh38 reference genome. The gene read quantification was performed using HTSeq-count (v0.11.2). The gene annotation used for quantification was Ensembl version 99. Using the gene read count matrix, CPM was calculated per sample on all annotated genes. EdgeR (v3.28.1) with TMM normalization was used to perform differential gene expression analysis. Benjamini and Hochberg FDR was computed to adjust p values obtained for differentially expressed genes. For pathway analysis, gene signatures were obtained from previous studies[41, 42] (Sup. Table 4). Thereafter, changes in gene expression were compared with gene signatures and enrichment scores were obtained (Sup. Table 5). The enrichment scores of gene signatures were estimated using R GSVA v1.36.2[43].

### Quantitative real-time PCR

Total RNA from cells was isolated using the NucleoSpin RNA II kit (740955, BIOKE) using the manufacturer’s instructions. Thereafter, 1μg of RNA from each sample was used to perform cDNA synthesis using the RevertAid First Strand cDNA synthesis kit (K1621, Thermo Fisher Scientific). Real time PCR was performed with GoTaq qPCR Master Mix (A6001, Promega) using CFX Connect Detection System (1855201, Bio-Rad). GAPDH was used as internal control for normalization. A list of primers that were used are provided in Sup. Table 6.

### MTS cell viability assay

To measure the proliferative capacities, cells were seeded on 96-well plates with 100μl of media. The following and subsequent days, 20μl of MTS solution was added per well and incubated in CO_2_ incubator for 1.5 hours. Thereafter, absorbance was measured on a luminometer at 490nm.

### Luciferase reporter assay

Luciferase reporter assays were performed using Dual luciferase reporter system (Promega). BRE-luciferase or NF-κB reporter plasmid was transfected with either control empty vector or TRAF4 and CMV-Renilla in HEK293T cells seeded on a 24-well plate. About 72 hours post-transfection, cells were stimulated overnight with BMP6 (50ng/ml) or TNF-α (10ng/ml) in serum free media. The following day, cells were lysed in Passive lysis buffer (Promega) and relative luciferase units and renilla values were measured using a Luminometer.

### Transwell migration assays

HT1376, T24 or MBT-2 cells were grown in 10 cm dishes and serum starved overnight. The following day, cells were trypsinized and resuspended in 0.5% serum containing media; about 50,000 were seeded onto the upper chambers. The lower chambers (wells) were filled with 2% serum containing media and incubated overnight. The following day, cells were fixed in ice-cold methanol for 10 minutes and stained with crystal violet solution. The inner side of the chambers were wiped clean using cotton swabs dipped in 1XPBS to remove remaining cells. Migrated cells were visualized through brightfield microscope and images were captured at four random sites and quantified.

### Site directed mutagenesis

PCR reactions were performed using Quik-Change XL kit by Agilent Technologies (catalogue no.200517-4) according to manufacturer’s instructions. The presence of mutants was confirmed by sequencing. The list of primers that were used is provided in Sup. Table 6.

### Immunoblotting

Cells were lysed in RIPA buffer (150mM NaCl, 1% Nonidet P-40, 0.5% sodium deoxycholate, 0.1% sodium dodecyl sulfate, 50mM Tris pH 8.0), supplemented with protease inhibitors and phosphatase inhibitors, 50mM sodium fluoride, 100mM β-glycerophosphate and 1mM sodium orthovanadate. Protein estimation was performed on the lysates and equal amounts of protein lysates were boiled in 2X sample buffer. Thereafter, samples were loaded onto 10% SDS-PAGE gels and transferred onto 0.45μM PVDF membranes (Millipore). The membranes were blocked in 5% milk and probed with specific antibodies overnight at 4°C. For visualization of protein signals, blots were incubated with secondary antibodies which were HRP-linked and detected using chemiluminescence. The following antibodies were used: TRAF4 1:2000 (D1N3A, CST), E-cadherin 1:1000 (Cat no. 610181, BD), N-cadherin 1:1000 (Cat no. 610920, BD), Vimentin 1:5000 (CST), SLUG 1:1000 (C19G7, CST), SNAIL 1:1000 (C15D3, CST), Flag 1:5000 (M2, Sigma Aldrich), phospho-Serine 1:1000 (612546, BD), SMURF1 1:1000 (45-K, Santa Cruz), Myc 1:5000 (9E10, Santa Cruz), HA 1:5000 (Y11, Santa Cruz), GFP 1:5000 (FL, Santa Cruz) and GAPDH 1:10,000 (MAB374, Millipore).

### Wound-healing assays on Incucyte

T24 cells were trypsinized and counted, then about 25,000 cells were seeded on each well of a 96-well plate (Essen ImageLock™) and let to attach in the CO_2_ incubator for 5 hours. Thereafter, media containing serum was removed and replaced with serum free media and cultured overnight. The following day, a woundmaker tool (4563, Essen) was used to produce wounds on the 96-well plate. After washing 2 times with 1XPBS, cells were replenished with 100μl of 0.5% serum containing media with the indicated treatments. The plate was then placed into the Incucyte Systems for Live-Cell Imaging and Analysis. Real-time images of (migrating) cells were captured every 1 hour and wound closure was analysed. About 10-12 well replicates were used for each condition to produce statistical error and significance.

### Modelling and simulation

The coordinates for PIP3 lipid was obtained from a crystal structure deposited in the Protein Data Bank (PDB ID: 4RWV). We considered only the truncated PIP3-diC4 derivative since the extended hydrocarbon tail was not critical for the modelling investigation. The phosphates in the lipid headgroup were considered to be completely deprotonated, resulting in a net charge of “−7e” on the lipid analogue. The trimeric state crystal structure of the TRAF domain from human TRAF4 was also downloaded from the Protein Data Bank (PDB ID: 3ZJB). PIP3-diC4 was docked into the structure using the program GOLD[44] by defining K313 (Chain A) and K345 (Chain C) as active site residues since both these lysines have been shown to be critical for interaction through mutational studies[16]. The pose of PIP3-diC4 which was most similar to the previously reported model by Rousseau *et al*[16] was chosen from the docking solutions. The lipid molecule was then symmetrically translated to attain a structural state with one PIP3-diC4 lipid bound at all the three interfaces formed between the monomers of the TRAF trimer. The S334 phosphorylated (pS334) state was subsequently modelled and the phosphate group in the residue was also considered to be completely deprotonated with a net charge of “−2e”. The amber library files for the phospho-serine residue was obtained from http://research.bmh.manchester.ac.uk/bryce/amber. The force field parameters with BCC atomic charges for PIP3-dic4 were derived from the GAFF2 database[45] through the Antechamber module in AMBER18 (Case, D.A., *et al* 2018, University of California, San Francisco).

Four different systems were subjected to molecular dynamics simulations. 1) TRAF (S334), 2) TRAF (pS334) 3) TRAF (S334): PIP3-diC4 and 4) TRAF (pS334): PIP3-diC4. The N- and C-termini of TRAF in all these systems were capped with ACE (acetyl) and NME (N-methyl) functional groups respectively. They were embedded in the centre of an octahedral box whose dimensions were fixed with a minimum distance of 8 Å between any solute atom and the box boundaries. The TIP3P water model was used for solvation. The net charge of all the systems was neutralized by adding appropriate number of counter ions. Molecular dynamics simulations were carried out using the PMEMD module in AMBER18. ff14SB[46] and GAFF2[45] force field parameters were used for the protein and PIP-diC4 respectively. All the systems were energy minimized, heated to 300 K over 30 ps by employing NVT ensemble and equilibrated for 200 ps in the NPT ensemble. The final production dynamics was run for 100 ns each under NPT conditions. A harmonic distance restrain was applied during the simulation between the phosphate atoms of the lipid (P3 and P5) and the side-chain nitrogen atom (NZ) of residues K313 and K345 with a force constant of 5 kcal/mol/Å^2^ [16] and an equilibrium distance of 4 Å. The simulation temperature of 300 K was maintained using Langevin dynamics with collision frequency of 1.0 ps^−1^. The simulation pressure was kept at 1 atm using weak-coupling with relaxation time of 1 ps. Periodic boundary conditions were applied and long-range electrostatic interactions was computed using Particle Mesh Ewald. The SHAKE algorithm was used to constrain all the bonds involving hydrogen atoms and the equation of motion was numerically solved with an integration time step of 2 fs.

### Immunohistochemical staining

Tissue microarrays containing bladder cancer samples of Stages 1, 2 and 3, as well as adjacent normal tissue and healthy bladder tissue were purchased from Biomax (BL802b, Biomax, U.S). Sections were heated at 60°C for 30 minutes prior to staining. Sections were deparaffinized and rehydrated, followed by heat induced antigen retrieval in 0.01M sodium citrate/0,05%Tween pH 6 NFκB-p65 (phospho-Ser311) and phospho-SMAD1/5/8 antibodies) or 10mM TRIS (pH9)/1mM EDTA/ 0,05% Tween (TRAF4 antibody) for 20 minutes. Sections were blocked for 30 minutes with 1%BSA and 0.1%Tween, followed by overnight incubation with primary antibodies at 4°C. Primary antibodies used were TRAF4 (1:50, HPA052377, Atlas, Bromma Sweden), NFκB-p65 (phospho-Ser311) (1:100, #11260, Signalway antibody, Uithoorn, the Netherlands) and phospho-SMAD1/5/8 (1:50, #9511, Cell Signaling, Leiden, The Netherlands). Sections were incubated with secondary antibody Alexa Fluor 488 donkey anti-rabbit (1:250, A-21206, Invitrogen, Landsmeer, the Netherlands) for 2 hours at room temperature, followed by 10 minutes of DAPI staining. Slides were digitalized using Pannoramic 250 flash III slide scanner (3DHISTECH, Budapest, Hungary) and staining for all antibodies were scored by two independent observers and their average scores were considered. TRAF4, NFκB-p65 (phospho-Ser311), and phospho-SMAD1/5/8s staining were scored combining the staining intensity (0: no staining, 1: low staining, 2: medium staining, 3: high staining) and percentage of positive tumour cells (TRAF4) or percentage of tumour cells with nuclear staining (NFκB-p65 (phospho-Ser311) and phospho-SMAD1/5/8 (0: 0%, 1: 1-5%, 2: 6-25%, 3: 26-50%, 4: 51-75%, 5: 76-100%). Staining from normal bladder tissue samples (n=8) were not considered for analysis for pSMAD1/5/8 and p-p65 samples. Representative photos of TRAF4 staining on grade 1, 2 and 3, as well as low and high NF-κB-phospho-p65 and phospho-SMAD1/5/8 staining were generated using the Caseviewer software version 2.0 (3DHISTECH, Budapest, Hungary).

### Statistics analyses

Bar graphs show mean standard deviation (SD) or mean standard error (SEM) as indicated in the figure legends. Student’s t test was used for the analysis of significance and p value. Kaplan-Meier graph was plotted using survival curve (GraphPad prism). For regression plots, Pearson’s r was used to analyse correlation. All tests were two-tailed.

## Author contributions

P.V.I., D.L.M., D.L., S.S., F.X., M.v.D., performed the experiments. P.V.I designed the experiments and interpreted the results. D.L. and C.S.V., performed the molecular modelling and simulations. T.Z.T. performed the EMT score, pathway analysis and gene signature studies. H.M performed the transcriptomic analysis of RNA-seq data. D.L.M. and L.R. performed the immunohistochemistry and scoring analysis. F.X. and L.Z. performed the proteomic studies. P.t.D. supervised the project. P.V.I. conceived the project and wrote the paper. All authors read, corrected and approved this manuscript.

## Acknowledgements

P.V.I thanks the support from the European Union’s Horizon 2020 research and innovation programme under the Marie Skłodowska-Curie Individual Fellowship. P.t.D and L.R. are supported by Cancer Genomics Centre, Netherlands. L.R. is also supported by a Veni grant (NWO, 016.176.081) and an LUMC Gisela Thier grant. We are grateful to Slobodan Vukicevics (University of Zagreb) for providing BMP6. We thank Midory Thorikay for technical assistance. Martijn Rabelink for shRNA constructs. MBT-2 cell line was a kind gift from Simon Dovedi. RNA-seq was performed by BGI Tech (Hong Kong). Karien Wiesmeijer and Annelies Boonzaier-van der Laan helped with scanning the tissue microarray slides. Also thanks to Kees Fluiter for providing advice on pathway studies.

## Figure legends

**Supplemental Figure 1**

**A** Immunoblot analysis probed with the indicated antibodies in cell lines treated with either control (DMSO) or 5’-Azacitydine, GAPDH shows loading control. **B** Multiple-sequence alignment indicating the conserved Serine-334 of TRAF4 in different species. **C** Immunoblot result from 293T cells transfected with either TRAF4 wildtype, S334A or S334E mutant; cells were treated with cycloheximide (CHX, 10μg/ml) for the indicated times and lysed. **D** Graph representing rate of degradation of proteins with respect to GAPDH. **E** Immunoblot result from 293T cells transfected with the indicated plasmids, MG132 was added overnight at 2μM concentration. **F** Immunoblot analysis from 293T cells transfected with the indicated plasmids, GAPDH shows loading control. **G** Immunoblot analysis from 293T cells transfected with the indicated plasmids, MG132 was added overnight at 2μM concentration where indicated, numbers represent relative quantification of TRAF4 levels. **H** Immunoblot analysis from 293T cells transfected with Myc-TRAF4; MG132 or MEK inhibitor (PD0325901) was added overnight at 2μM concentration as indicated. Numbers represent relative quantification of TRAF4 levels with respect to GADPH. **I** Immunoblot analysis from UMUC3 cells, treated overnight with MEKi at 2μM concentration as indicated, numbers represent relative levels of TRAF4. **J** Real-time PCR analysis of *TRAF4* mRNA expression levels after treating the cells with MEKi overnight as indicated; error bars represent ± SD, *n.s*. denotes non-significant P value calculated using two-tailed student’s t test. **K** Immunoblot analysis from T24 cells, treated overnight with MEKi at 2μM concentration as indicated, numbers represent relative levels of TRAF4 with respect to GAPDH. **L** Real-time PCR analysis of *TRAF4* mRNA expression levels after treating the cells with MEKi overnight as indicated, *n.s*. denotes non-significant P value.

**Supplemental Figure 2**

**A** Relative fluorescence intensity of points across the plasma membrane transfected with either GFP, GFP-TRAF4, GFP-TRAF4 mutants (S334E and S334A), peaks indicate strong positive signal across the membrane. **B** HT1376 cells transfected with GFP vector. **(C and D)** Representative structures from the molecular dynamics simulations of the S334 and pS334 structural states of the TRAF trimer. **(E and F)** Magnified image of one of the interfacial regions involved in the recognition of PIP3 from the S334 and pS334 structures. **(G and H)** Magnified image of one of the interfaces from the S334 and pS334 structures in the presence of PIP3-diC4 lipid. See Figure 2I in the main text for the full trimer representation. The protein in all the figures is shown in electrostatic surface representation which was created using the APBS plugin through the Pymol molecular visualization software (Schrondinger). The color gradient from blue to red indicates the range of electrostatic surface potential kT/e values from strongly positive (+3.0) to strogly negative (−3.0). The PIP3-diC4 lipid and specific protein residues which are involved in the interaction with it’s headgroup are shown in stick representation.

**Supplemental Figure 3**

**A** Immunoblot result showing TRAF4 levels in RT4 cells stably expressing short hairpins to either control (empty vector pLKO) or TRAF4. **B** Wildtype RT4 cells (a colony) stained with CellMask™ Orange plasma membrane stain and image captured through fluorescence microscopy. **C** Immunoblot result showing TRAF4 levels in HT1376 cells stably expressing short hairpins to either control (empty vector pLKO) or TRAF4. **D** Real-time PCR result showing mRNA expression levels of *SNAI1* in bladder cancer cell lines. **E** Real-time PCR result showing mRNA expression levels of *SNAI2* in bladder cancer cell lines.

**Supplemental Figure 4**

**A** Immunoblot analysis showing TRAF4 and other EMT markers protein expression in bladder cancer cell lines including MBT-2. **B** Immunoblot result showing Smurf1 levels in MBT-2 cells stably expressing short hairpins to either control (empty vector pLKO) or *SMURF1*. **C** Brightfield images of MBT-2 (control and TRAF4 over-expressing) cells seeded and grown at low density, scale: 400μM (representative images). **D** Immunoblot showing T24 stably expressing TRAF4, A-D are the four replicates used for transcriptomic analysis. **E** Graph represents changes in enrichment scores of eleven major cancer associated signalling pathways in T24 cells when TRAF4 is stably over-expressed.

**Supplemental Figure 5**

**A** Real-time PCR result showing *ID3* mRNA expression levels in BMP6 (50ng/ml for 2 hours) treated MBT-2 cells as indicated; error bars represent ± SD, \**P* ≤ 0.05 calculated using two-tailed student’s t test. **B** Luciferase reporter assay in HEK293T transfected with BRE-luciferase reporter, SV40 Renilla and either empty vector control or TRAF4, transfected cells were stimulated overnight with BMP6 (50ng/ml) where indicated, error bars represent ± SD, \*\*\**P* ≤ 0.001, calculated using two-tailed student’s t test. **C** Graph showing relative wound density result using T24 cells from Incucyte, TRAF4 over-expressing cells were treated with a BMP receptors inhibitor LDN193189 at 120nM concentration where indicated. Images were obtained every 1 hour after wound was produced; error bars represent ± SEM. **D** Representative images from Incucyte result, brown area represents the cell coverage and grey area is the wound produced and remaining. **E** Luciferase assay in 293T transfected with NF-κB luciferase reporter, SV40 Renilla and either empty vector control or TRAF4, transfected cells were stimulated overnight with TNF-α (10ng/ml) where indicated, error bars represent ± SD, \**P* ≤ 0.05, calculated using two-tailed student’s t test. Representative result from three independent experiments.

