## Supplemental figures for "TRAF4 inhibits bladder cancer progression by promoting BMP/SMAD signalling pathway"

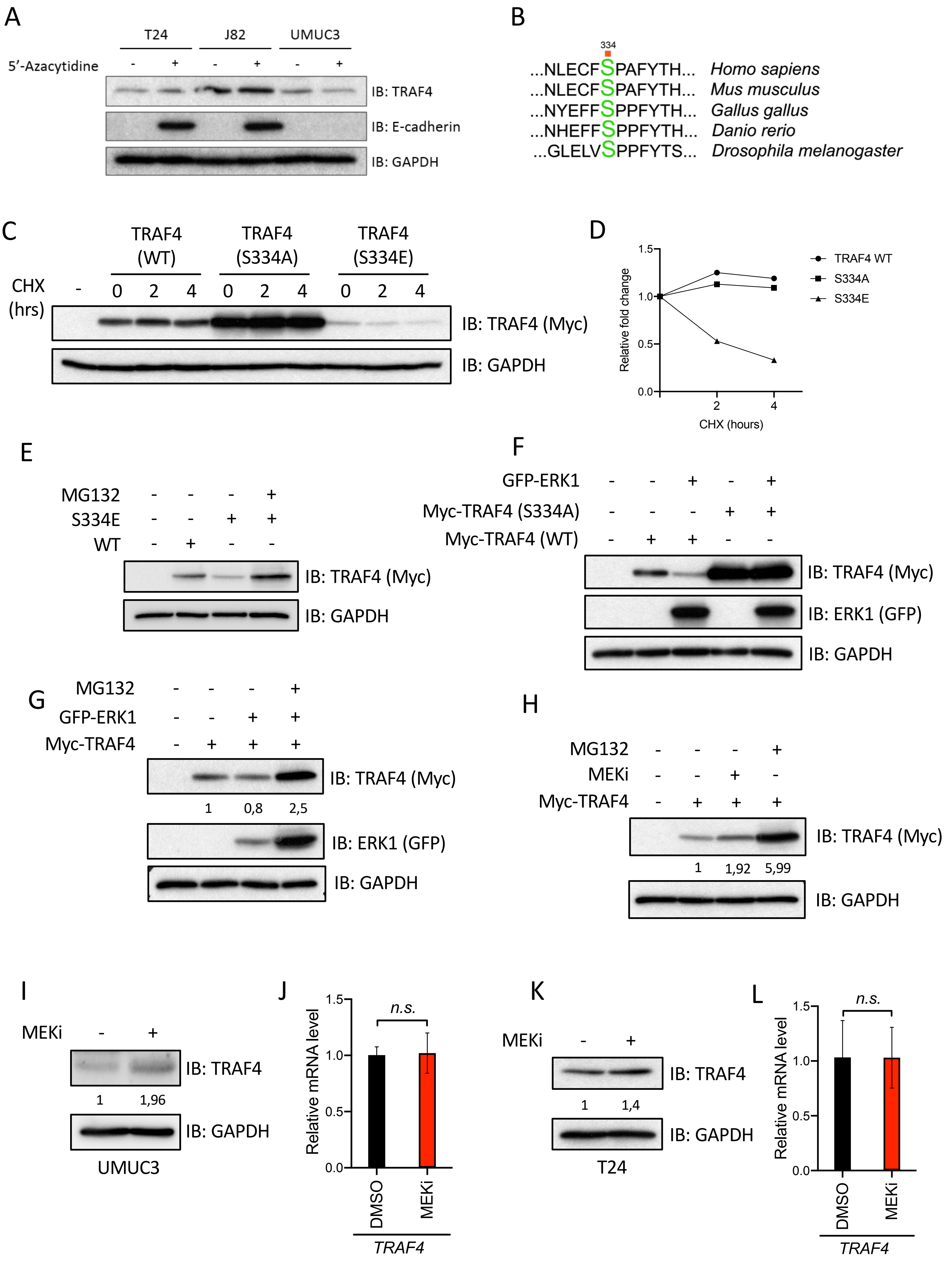


Supplemental Figure 1

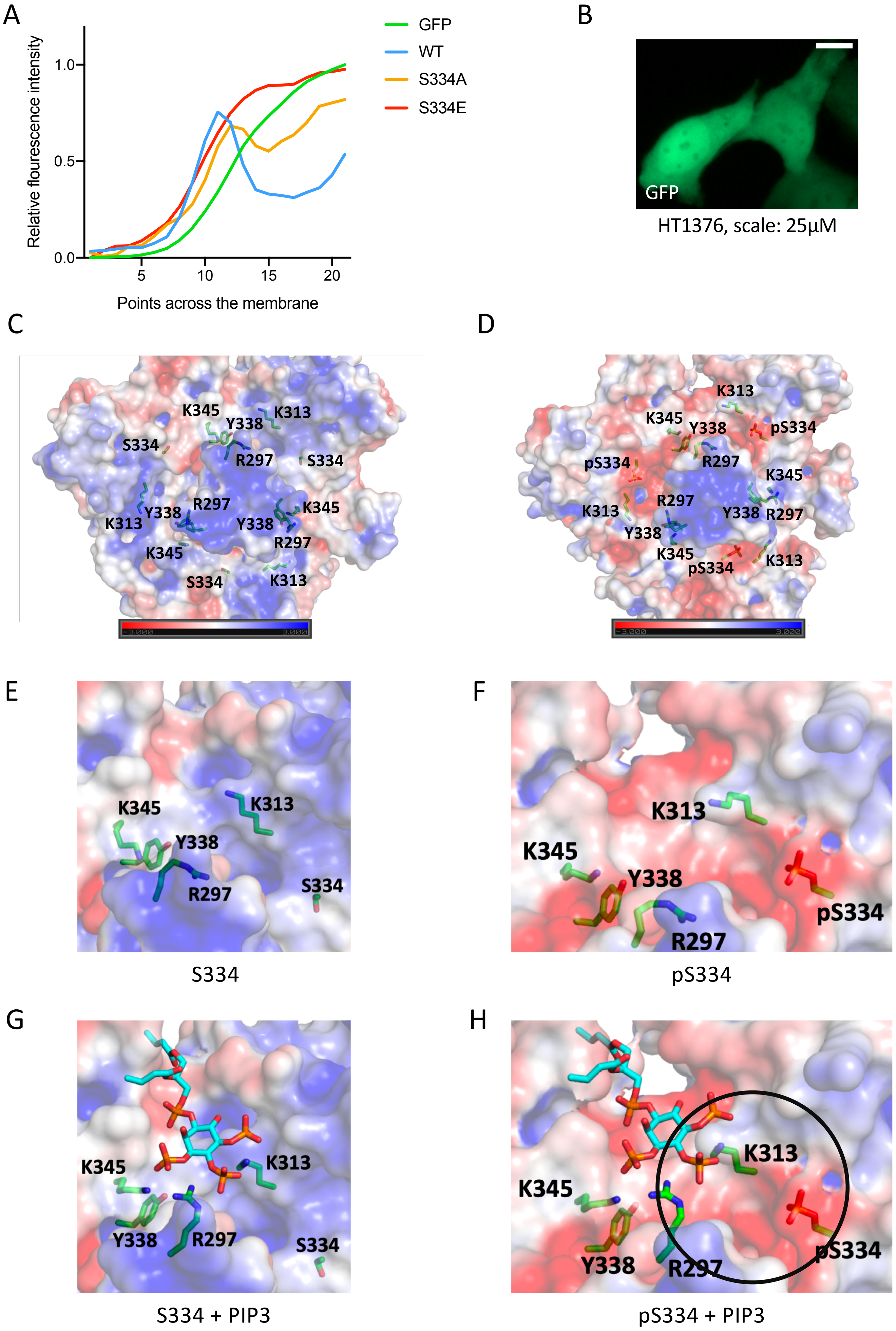


Supplemental Figure 2

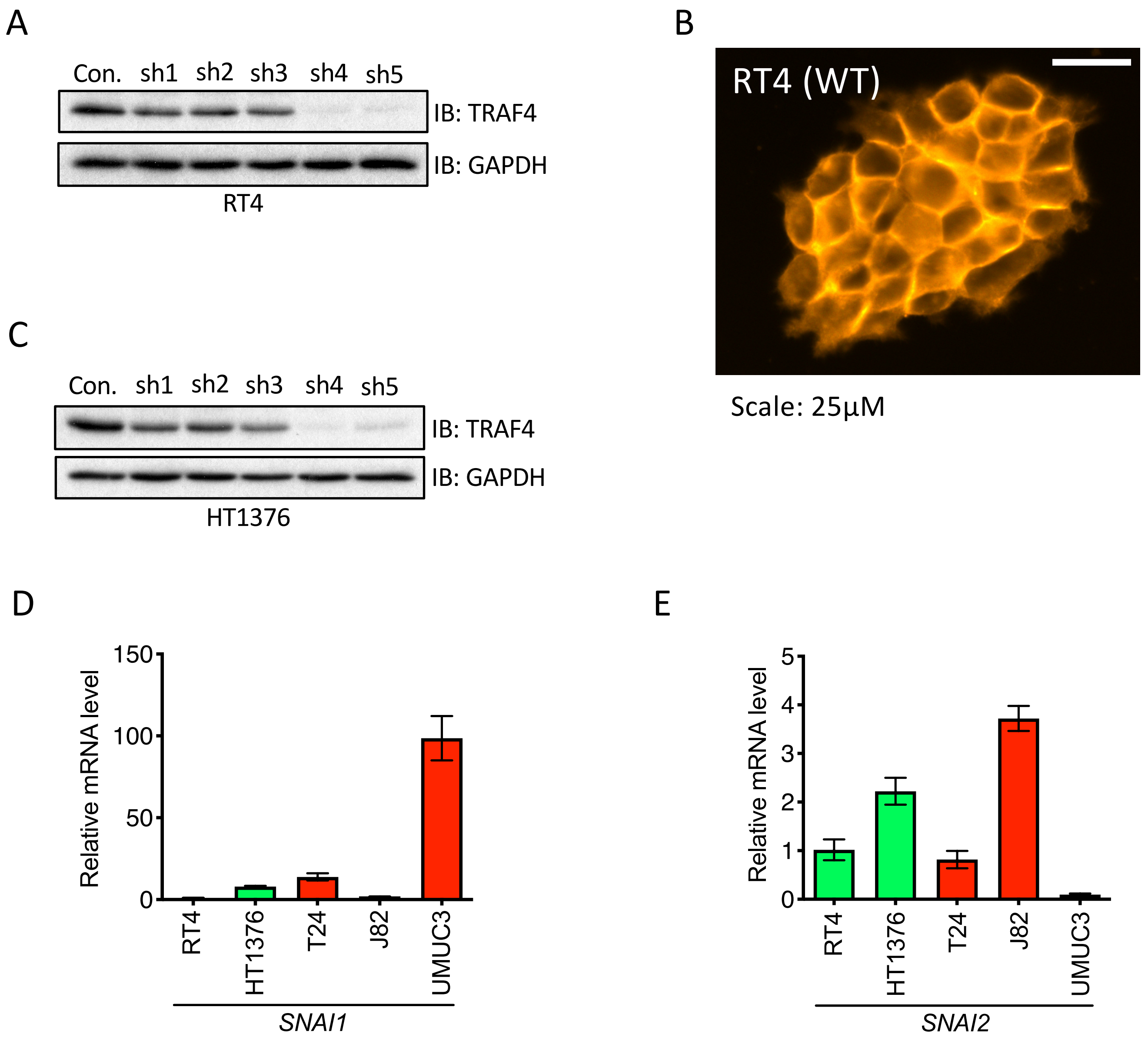


Supplemental Figure 3

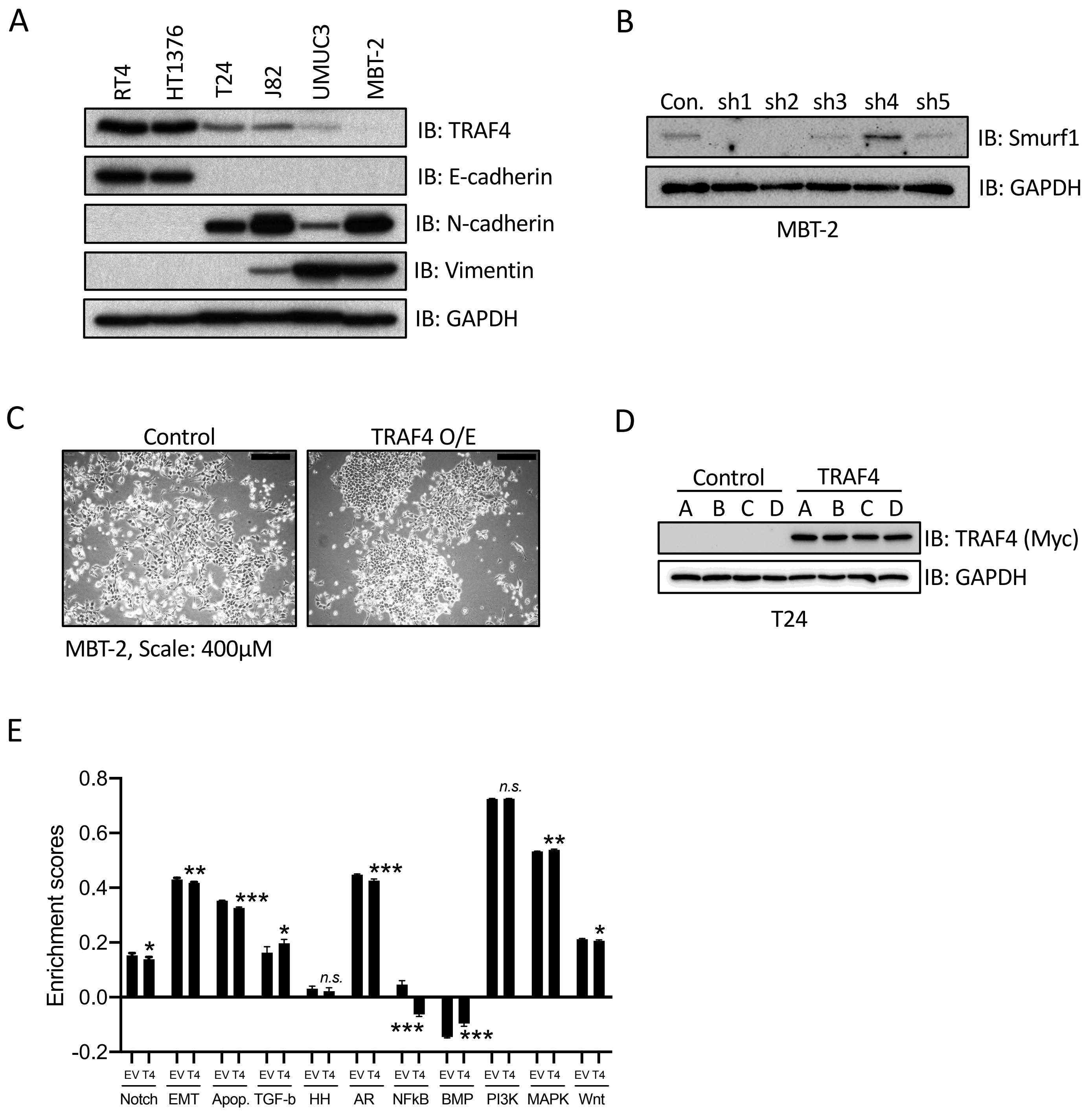


Supplemental Figure 4

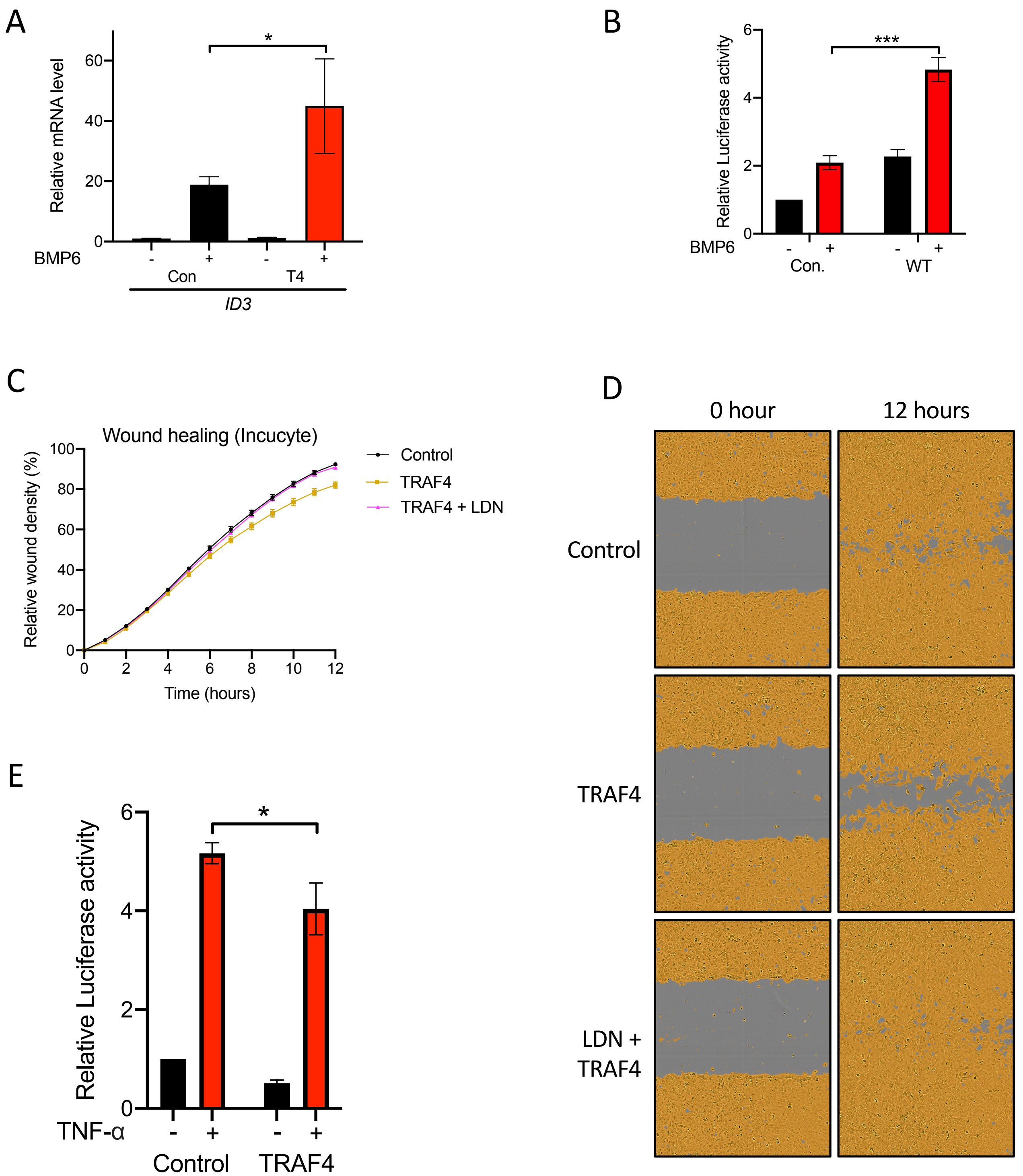


Supplemental Figure 5
